# Regenerative potential varies along the anterior-posterior axis of the annelid *Capitella teleta*

**DOI:** 10.64898/2025.12.19.695534

**Authors:** Lauren F. Kunselman, Elaine C. Seaver

## Abstract

Regeneration abilities vary among species, but they can also vary within individuals depending on factors such as developmental stage, amputation location and nutritional status. Many annelids (segmented worms) have well-documented cases of regeneration variation along the anterior-posterior (AP) body axis. However, mechanistic explanations for the diverse regeneration outcomes at different amputation positions remain unknown. *Capitella teleta* is an annelid with robust posterior regeneration ability but lacks anterior regeneration, although anterior regeneration ability has not been rigorously assessed at multiple amputation sites. In this study, we characterize regeneration ability along the AP axis of *C. teleta* tail fragments by comparing the post-amputation response of tail fragments cut between segments 1 and 2 and tail fragments cut between segments 10 and 11. Through EdU experiments, *in situ* hybridization assays, and antibody labeling, we demonstrate that the more anterior amputation site proceeds to later stages of regeneration than the posterior amputation site, but regeneration does not go to completion. The distribution of neoblast-like cells after amputation suggests that these cells do not substantially contribute to formation of the anterior-facing blastema in tail fragments with higher inherent regeneration potential. Lastly, we test whether the greater regenerative competence of tail fragments amputated between segments 1 and 2 increases the probability of accomplishing complete posterior regeneration after treatment with CHIR, a Wnt/β-catenin agonist. Regeneration outcomes are comparable following increase in Wnt/β-catenin signaling regardless of amputation position, suggesting that initial regeneration potential is not a limiting factor of successful posterior regeneration. Comparing tissues with different regenerative abilities within an individual organism can elucidate mechanisms underlying regeneration regulation, thereby enabling the prospect of rescuing or increasing regeneration ability in regeneration-deficient tissues.

## Introduction

Regenerative abilities vary widely among animals, and interspecies comparisons of regeneration ability offer the promise of mechanistic explanations for naturally occurring regeneration failure and limitations in many species and lineages (Bely, 2010). In addition to interspecies comparisons of regeneration abilities, there is another important aspect of regeneration that could contribute to our understanding of regeneration constraints, which is only recently gaining more attention. Namely, regeneration ability can also vary within a single individual. For example, although planarians are best known for regenerating a complete animal from many different amputation planes made along the body, there are some species of planarians that fail to regenerate a head if they are cut transversely in the posterior third of their body (Liu et al., 2013; Sikes & Newmark, 2013). Similarly, although many species within the *Hydra* genus robustly regenerate their whole body, *Hydra oligactis* fails to regenerate a foot with a higher frequency when it is cut closer to the head (Campos et al., 2025).

Numerous descriptions of intraindividual variation in regeneration ability come from annelid regeneration research. This is likely because the segmented body plan that characterizes the phylum makes them amenable to precise cuts along their anterior-posterior (AP) axis, and furthermore, members of this phylum display a wide variety of regeneration abilities, ranging from inability to regenerate to whole-body regeneration from a single segment (Bely, 2006; Seaver & de Jong, 2021). For example, *Chaetopterus variopedatus* can regenerate a head from any segment anterior to segment 14, but head regeneration fails when amputations are made posterior to that segment (Berrill, 1952). In *Enchytraeus buchholzi*, the number of head segments regenerated decrease as amputations are made further from the head, and when amputations are made close to the tail, an ectopic tail is regenerated, forming a bicaudal animal. A similar phenomenon occurs in *Lumbriculus inconstans* (Hyman, 1916). When *Enchytraeus japonensis* is amputated within the seven anterior-most segments that make up the head region, the number of head segments that regenerate depend upon the axial position of the cut (Myohara, 2012). However, *E. japonensis* has no variation in anterior or posterior regeneration ability throughout its trunk region (Myohara, 2012). *Platynereis dumerilii* tail regeneration occurs at a faster rate from more anterior cuts, until an amputation is made within one segment of the pharynx, upon which it loses the ability to regenerate a tail (Planques et al., 2019).

Despite the plethora of examples of varying intraindividual regeneration ability in many annelid species, mechanistic explanations for these diverse regenerative responses are lacking. Various physiological hypotheses have been put forward. For instance, when tissues or structures in a particular region of the body inhibit proper wound healing following injury, regeneration often fails and the injury leads to mortality (Planques et al., 2019; Spieß et al., 2024). For example, amputations near the pharynx of *P. dumerilii* lead to eversion of the pharynx, failed wound closure, and subsequent death (Planques et al., 2019). Additionally, a small fragment of tissue remaining after an injury may have less regeneration potential because it cannot meet the energetic demands required to grow new tissue (Dirks et al., 2012; Hyman, 1916). These processes do not satisfactorily explain all cases of variable intraindividual regeneration ability, however.

Another plausible mechanism for regeneration failure along the AP axis could be related to the distribution and/or behavior of stem cells during regeneration. Stem cells are reported to serve as the source of regenerated tissue in several annelids, although local cellular sources have been implicated in annelid regeneration as well (de Jong & Seaver, 2018; Kostyuchenko & Kozin, 2020; Myohara, 2012; Novikova et al., 2013; Planques et al., 2019; Randolph, 1892; Tadokoro et al., 2006; Takeo et al., 2008; Tanaka & Reddien, 2011). In some organismal models of stem cell-based regeneration, if an amputation is made and a resulting tissue fragment lacks stem cells, regeneration can fail. For example, the pharynx of the planarian *Schmidtea mediterranea* lacks neoblasts, and when isolated from the animal it cannot regenerate a complete animal, although a tissue fragment of comparable size containing neoblasts can regenerate (Rossant, 2014). A lack of stem cells in the head portion of the body has been proposed as the underlying reason for the restricted anterior regeneration ability of *E. japonensis* in this region; an interpretation based also on its consistent regeneration ability throughout the trunk region, which contains stem cells (Myohara, 2012). This hypothesis has not been functionally tested, however, by say, selectively ablating stem cells in the trunk or transplanting stem cells to the head and then assessing regeneration outcome after either procedure.

Another common feature of stem cell-based regeneration models is that stem cells utilize guidance cues to migrate towards the wound site so they can serve as the cellular source for new tissue (Bradshaw et al., 2015; Guedelhoefer IV. & Alvarado, 2012; Ullah et al., 2019). *In vivo* migration of stem cells towards the wound has been observed in decapitated transgenic *Hydractinia echinata* (Bradshaw et al., 2015). Through tissue transplantation and partial irradiation, directional migration of stem cells towards anterior or posterior amputation sites has also been demonstrated in *S. mediterranea* (Guedelhoefer IV. & Alvarado, 2012). Cells with neoblast-like qualities have been observed migrating towards anterior and posterior wounds in the annelid *Pristina leidyii*, but the identity of the cells have not yet been verified (Zattara et al., 2016). If there are limitations on where stem cells can migrate due to failure to recognize and respond to signals from the wound or morphological obstructions between the stem cell and the wound, this could also explain variation in regeneration potential within an individual. However, there is still a general lack of understanding for how stem cell behavior may influence regeneration variation within individuals of a species.

Annelids have immense unrealized potential to offer novel insights into the causes of regeneration constraints. To pioneer this avenue of research, we turned to a well-established annelid regeneration model, *Capitella teleta*. *C. teleta* exhibits an interesting mixture of both robust regeneration abilities and some regeneration limitations that make it amenable for many comparisons of regeneration regulation within an individual. *C. teleta* can regenerate a tail from almost any position along its AP axis (de Jong & Seaver, 2016), but this ability declines as amputations are made within the anterior-most six segments of the thorax in juveniles (LK, personal observations). Head regeneration is not observed in *C. teleta* (Bely, 2006; Kunselman & Seaver, 2025), although this has not been assessed in depth at multiple amputation positions along the AP axis. *C. teleta* also seems to utilize different sources of cells when regenerating tissue. Experimental evidence suggests a putative cluster of multipotent progenitor cells, referred to as the MPC cluster, is a source of migrating stem cells that contribute to posterior regeneration in *C. teleta* (de Jong & Seaver, 2018). Additionally, EdU-pulse-chase experiments suggest that dividing cells proximal to the amputation site contribute to the blastema (de Jong & Seaver, 2018). The perfect balance of regeneration capabilities and regeneration constraints, combined with different methods of regenerating new tissue, make *C. teleta* an excellent model to begin to answer the question of how regeneration ability is regulated in different cell types and tissues within an individual.

In this study, we investigate the regeneration potential along the AP axis of *C. teleta* tail fragments by comparing the post-amputation response of tail fragments cut at multiple amputation locations. Through EdU experiments, *in situ* hybridization assays, and antibody labeling, we find that regeneration progresses to later stages at anterior amputation sites relative to more posterior amputation sites, but regeneration does not go to completion. We also explore whether the MPC cells contribute to the increased regeneration potential of the more anterior amputation site by observing their distribution following amputation. Our data suggest that they are not a major source of cells for the anterior-facing blastema. Finally, we test whether the greater regenerative competence of tail fragments amputated between segments 1 and 2 increases the likelihood of achieving complete posterior regeneration after treatment with CHIR, a Wnt/β-catenin agonist. We find that similar regeneration outcomes occur regardless of amputation position, suggesting that initial regeneration potential is not a limiting factor of successful posterior regeneration. Our results raise compelling questions about the underlying causes of variable intraindividual regeneration potential and mechanisms of blastema formation at different locations within an organism. An understanding of the regulatory feedback governing tissue regeneration could unlock latent or repressed regeneration potential. Our ability to harness this process depends on our comprehension of its underlying cellular and molecular mechanisms, and *C. teleta* and other annelids are fitting research models to lead us closer to this goal.

## Methods

### Animal Husbandry and Amputations

A *C. teleta* colony was maintained in the laboratory at 19°C according to published culture methods (Grassle & Grassle, 1976). Animals were fed previously frozen and sieved estuarine mud and fresh sea water once per week. Naturally developed late-stage larvae from the colony were transferred to bowls of mud and filtered seawater (FSW) and raised for two-weeks post-metamorphosis as juvenile worms for regeneration experiments.

For amputations, two-week-old juvenile worms were removed from the mud and allowed to burrow in a 35 mm petri dish filled with 0.5 % cornmeal agar (Sigma-Aldrich) in FSW for 2-5 hours to clear their gut contents and to remove debris from the outside of the worm. Then, worms were immobilized in 1:1 FSW:0.37M MgCl_2_ for approximately 15 minutes, placed in a drop of FSW:MgCl_2_ solution on an amputation platform of black dissecting wax (American Education Products), and using a dissection microscope (Stemi 2000, Zeiss) for visualization, amputated with a microscalpel (Feather, 15° blade). Tail fragments cut between segment 10 and 11 (10-11 amputees) were placed in FSW supplemented by 60 µg/mL penicillin plus 50 µg/mL streptomycin, and water was changed approximately every three days. Tail fragments cut between segment 1 and 2 (1-2 amputees) or 4 and 5 (4-5 amputees) were placed in 1:4 0.37M MgCl_2_:FSW supplemented by 60 µg/mL penicillin plus 50 µg/mL streptomycin that was changed daily. This gentle anesthetic kept the longer tail fragments from adhering to the bottom of the dish with mucus or tangling up with other fragments in the dish and resulted in improved viability. Tail fragments were monitored daily to remove any mucus or gut contents they secreted over time. Tail fragments are unable to feed; therefore, no mud was provided.

### Whole Mount *in situ* Hybridization

Prior to fixation, animals were cleaned of sediment, if necessary, by placing them in a cornmeal agar plate (see above) for 2-4 hours. Then, they were exposed to a solution of 1:1 FSW:0.37M MgCl_2_ for 15 minutes and fixed in 4% paraformaldehyde (PFA) in FSW at 4°C for at least 24 hours. After fixation, intact and regenerating juveniles were washed in phosphate buffered saline (PBS) + 0.2% Triton (PBT), dehydrated in a methanol series to 100% methanol, and stored at −20°C. Anti-sense digoxigenin-labeled (Sigma-Aldrich) riboprobes for all genes were synthesized in vitro with the SP6 or T7 MEGAscript kit (Ambion, Inc., Austin, TX, USA). Probe lengths are as follows: *CapI-vasa* (NCBI accession number BK006523): 1122 bp, 1.5 ng/µL; *Ct-NeuroD* (NCBI accession number MF508644): 669 bp, 1 ng/µL; *Ct-otx*: 1030 bp, 0.2 ng/µL. Riboprobes were diluted with hybridization buffer to a concentration of 50 ng/µL and stored at −20°C. Genes published prior to the formal species designation of *C. teleta* (Blake et al., 2009) use *CapI* as an abbreviation for *Capitella sp. I* in their naming, and they are cloned from the same species as genes with the *Ct-* (*C. teleta*) prefix.

Whole mount *in situ* hybridization followed published protocols (Seaver & Kaneshige, 2006). After hybridization at 65°C for 72 hours followed by an overnight exposure at 4°C to an anti-digoxigenin antibody (Roche Diagnostics), probes were visualized with nitro blue tetrazolium (Thermo Scientific)/5-bromo-4-chloro-3-indolyphoshate (US Biological) (NBT/BCIP) color substrate. The duration of the colorimetric reaction was typically between 1 hour and 3 days. For longer development times, specimens were usually incubated with substrate at room temperature during the day and at 4°C overnight until development was complete. DMSO controls were developed for the same amount of time and temperature as their CHIR-treated counterparts within each experiment. In cases of high background or crystal formation during the color reaction, head and tail fragments were washed following termination of the colorimetric reaction by sequential ethanol clearing steps (Siebert et al., 2019). This comprised a 5-minute wash in 33% ethanol (in PBT), a 5-minute wash in 66% ethanol (in milliQ water), and incubation in 100% ethanol until precipitate appeared blue (usually 30 minutes). Finally, samples were rehydrated in 66% ethanol (in milliQ water) for 5 min, followed by a 5-minute wash in 33% ethanol (in PBS), a 5-minute wash in PBT, and then transferred to PBS.

Samples were mounted in 80% glycerol in PBS and placed on Rainex-coated slides prior to microscopic analysis. At least 2 independent trials were performed for each gene.

### Detection and Quantification of Cell Proliferation

The Click-iT EdU Alexa Fluor 488 Imaging Kit (Invitrogen) was used to label cells in the S-phase of the cell cycle according to manufacturer instructions. Tail fragments were exposed to 5’-ethynyl-2’-deoxyuridine (EdU) at a final concentration of 3 µM for 1 hour at either 3 dpa or 7 dpa. Animals were then placed in 1:1 FSW:0.37MgCl_2_ for 15 minutes followed by fixation. For specimens intended to be labelled with the anti-Vasa antibody, animals were fixed in 0.5% PFA:FSW for 1 hour at RT followed by a 5 minute wash in 0.5% SDS. Otherwise, animals were fixed in 4% PFA:FSW for 30 minutes – 1 hour at RT. After performing the EdU detection reaction, samples were subjected to immunohistochemistry.

To quantify cell proliferation, confocal z-stack images were cropped to the area of interest– defined as the segment closest to the amputation site plus any new tissue growth following amputation. A segment was defined as the anterior edge of a ganglion to the anterior edge of the ganglion in the adjacent segment. EdU-positive nuclei and total nuclei (counter-stained with Hoechst 33342 (Molecular Probes)) were quantified using Imaris Software (V7.6.1, Bitplane, Switzerland) using a size threshold of 2.55 µm and a quality score of >10. Each area of interest for each specimen was manually inspected to ensure that digital identification of nuclei was accurate. The number of EdU-positive nuclei was divided by the number of total nuclei and multiplied by 100 to generate a percentage for each sample. A Wilcoxon rank sum test was used to determine if there was a significant difference in the average percentage of EdU-positive cells between two treatments at the same time point. Differences in averages with a p value <0.05 were considered statistically significant.

### Immunohistochemistry

Prior to fixation, animals were cleaned of sediment, if necessary, by placing them in a cornmeal agar plate for 2-4 hours. Then, animals were exposed to a solution of 1:1 FSW:0.37M MgCl_2_ for 15 minutes and fixed in 4% PFA in FSW at room temperature (RT) for 30 minutes - 1 hour. Following fixation, tissue fragments were washed in PBT several times and then placed in a blocking solution (PBT + 10% normal goat serum, Sigma G9023) with rocking for 1 hour. Primary antibodies were diluted in block solution and animals were incubated overnight at 4°C. Primary antibodies and the working dilutions used are as follows: 1:400 mouse anti-acetylated α-tubulin (Sigma T6743), 1:10 mouse anti-Pax (monoclonal antibody clone DP311, gift of Gregory Davis and Nipam Patel; Davis, 2002; Webster et al., 2021), 1:400 rabbit anti-FMRFamide (Immunostar, cat. 20091), 1:200 rabbit anti-serotonin (5HT, Sigma-Aldrich, cat. S5545), and 1:500 rabbit anti-Vasa (Bethyl Laboratories). The anti-Vasa antibody is a custom anti-peptide polyclonal antibody and its labeling pattern closely resembles published *in situ* hybridization patterns for *CapI-vasa* (Dannenberg & Seaver, 2018; de Jong & Seaver, 2018; Dill & Seaver, 2008). Animals were then washed twice in PBT followed by four 30-minute washes in PBT. Specimens were incubated overnight at 4°C in secondary antibody diluted 1:400 in blocking buffer. Secondary antibodies are as follows: goat anti-mouse Alexa Fluor 488 (Invitrogen), goat anti-mouse Alexa Fluor 594 (Invitrogen), goat anti-rabbit Alexa Fluor 488 (Invitrogen), goat anti-rabbit Alexa Fluor 594 (Invitrogen), and goat anti-rabbit Alexa Fluor 647 (Invitrogen). Secondary antibodies were washed from tissue fragments with the same number of PBT washes as when the primary antibody was removed. Hoechst 33342 (1mg/mL, Molecular Probes) was diluted 1:1000 in PBT and added to the last two 30-minute washes to label nuclei. Specimens were mounted in 80% glycerol:PBS on glass slides and then imaged and analyzed as described below.

### Pharmacological Activation of Wnt/β-catenin Signaling

To mimic activation of the Wnt signaling pathway CHIR-98014 (Millipore), an inhibitor of GSK3-β, was diluted to 5 µM in FSW from a 1 mM working stock solubilized in dimethyl sulfoxide (DMSO). CHIR was diluted in 1:4 0.37M MgCl_2_:FSW supplemented by 60 µg/mL penicillin plus 50 µg/mL streptomycin for 1-2 and 4-5 cut tail fragments to increase their survival, as described in an earlier section. Approximately 20 tail fragments were added per well to a 35 mm petri dish in 2 mL of CHIR solution, or 0.5% DMSO as a negative control. Animals were transferred daily to fresh CHIR or DMSO solution for the duration of the experiment. The CHIR concentration for the experiments was chosen based on previously published work (Kunselman & Seaver, 2025).

### Microscopy and Imaging

A Zeiss LSM 710 confocal microscope (Zeiss, Gottingen, Germany) coupled with a camera was used to image immunolabelled animals and EdU-treated samples. Images were captured with Zen Software (Zen 2011, V14.0.17.201, Zeiss, Gottingen, Germany). Z-stack maximum-intensity projections were generated using ImageJ (NIH).

To image specimens processed for *in situ* hybridization, an Axioscope 2 mot-plus compound microscope (Zeiss, Gottingen, Germany) was fitted with a SPOT FLEX digital camera (Diagnostic Instruments, Inc., Sterling Heights, MI) and images were captured with SPOT software. For some images, several DIC focal planes were merged using Helicon Focus (Helicon Soft Ltd., Kharkov, Ukraine). Images were processed using Adobe Photoshop 2024. Any adjustments of brightness or contrast made in Photoshop were applied to whole images. Figures were created using Adobe Illustrator 2024.

## Results

The segmented body of *C. teleta* juveniles facilitates precise amputations along the AP axis (Fig. 1a) and we leveraged this feature to compare anterior regeneration potential at two distinct axial positions. Tail fragments amputated between segment 10 and 11 (referred to as 10-11 amputees) and tail fragments amputated between segment 1 and 2 (referred to as 1-2 amputees) were compared over the course of a week following amputation (Fig. 1b-d). To explore the contribution of the MPC cluster to regeneration in anterior-facing wounds, we later performed transverse amputations at an additional location, between segment 4 and 5 (Fig. 1e).

**Figure 1.**
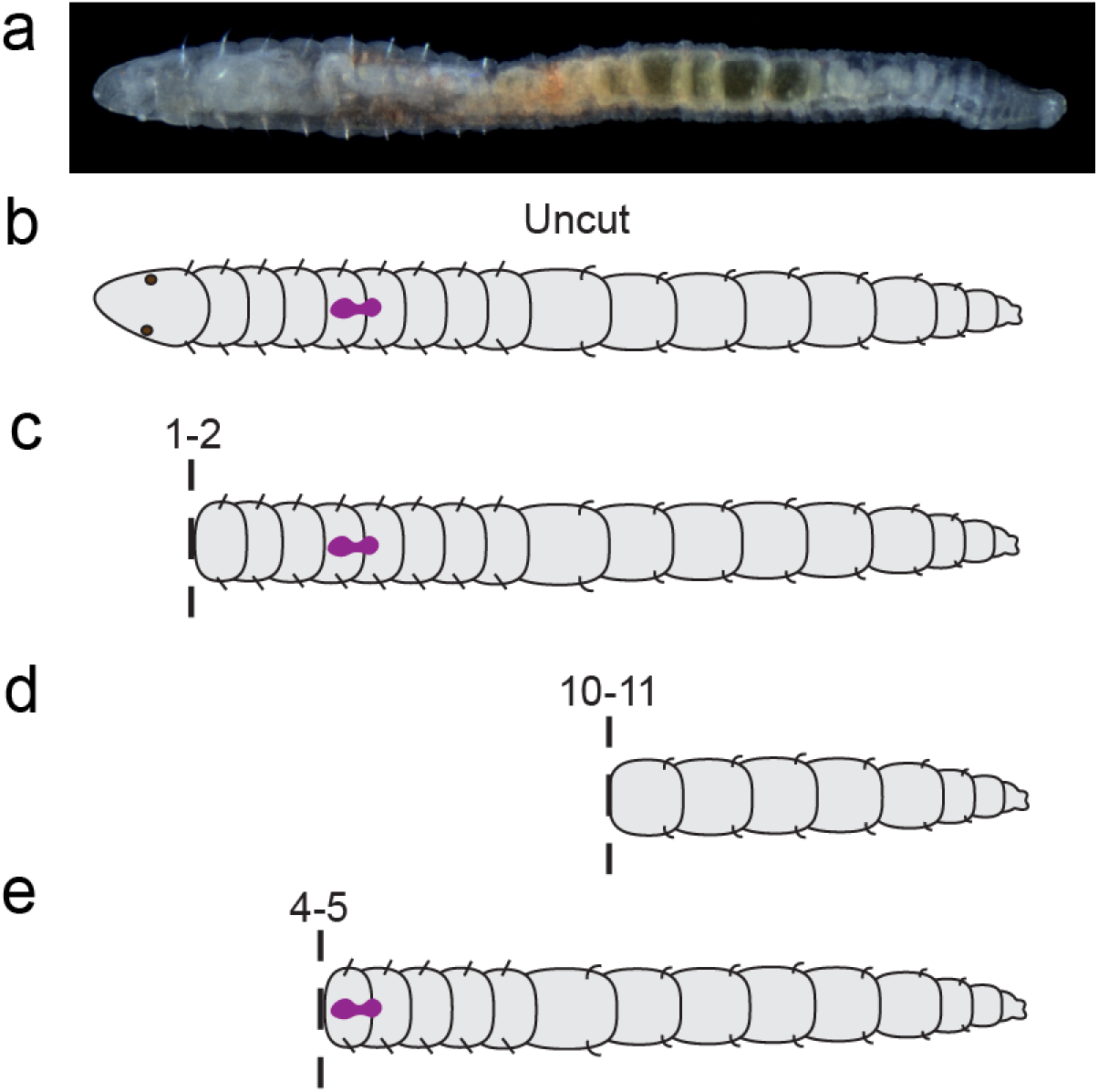
Amputation sites investigated along the AP axis of *C. teleta*. All images and graphics are in ventral view, and anterior is to the left. a) DIC image of an intact *C. teleta* juvenile. b) Graphic of intact *C. teleta*. c) Graphic of a tail fragment cut between segments 1 and 2. d) Graphic of a tail fragment cut between segments 10 and 11. e) Graphic of a tail fragment cut between segments 4 and 5. In b, c and e, the MPC cluster is represented by the purple hourglass shape. Dotted line indicates amputation plane.

### Differences in Regeneration Potential at Two Positions Along the AP Axis

The first step of regeneration, wound healing, occurs in all tail fragments regardless of amputation position, either by constriction of muscles to seal the wound or by extrusion of the gut to form a plug (data not shown). A subsequent early step of regeneration in *C. teleta* is when a blastema consisting of proliferating cells appears at 3 dpa (de Jong & Seaver, 2016). EdU labeling of cells in the S-phase of the cell cycle reveals localized cell division at the cut site in 1-2 amputees (Fig. 2a). In contrast, 10-11 amputees have sparse EdU-positive nuclei at the cut site 3 dpa, in accordance with previous results (Fig. 2b) (Kunselman & Seaver, 2025). The percentage of dividing cells is significantly higher in 1-2 amputees compared to 10-11 amputees, suggesting that the more anterior amputation position elicits more cell division (p= 2.0e-6, Wilcoxon rank sum test, Fig. 2c). Although these results are intriguing, it is important to note that in the uncut animal, EdU patterns are not distributed uniformly across the body. The pharynx, whose anterior edge is at the border of segments 1 and 2, is a tissue that exhibits high amounts of cell proliferation in uncut animals (Fig. S1a-d, brackets). The boundary of segments 10 and 11 typically has some cell division spread mostly across the ectoderm (Fig. S1e-h). Since cell division is abundant in the pharynx, there is the possibility that the higher amount of cell division seen at the wound of 1-2 amputees may be due to endogenous cell division characteristics of different tissues along the AP axis rather than initiation of a regenerative response. However, most of the EdU observed in 1-2 amputees is distal to the amputation site and therefore unlikely to be part of the intrinsic proliferation pattern in the pharynx.

**Figure 2.**
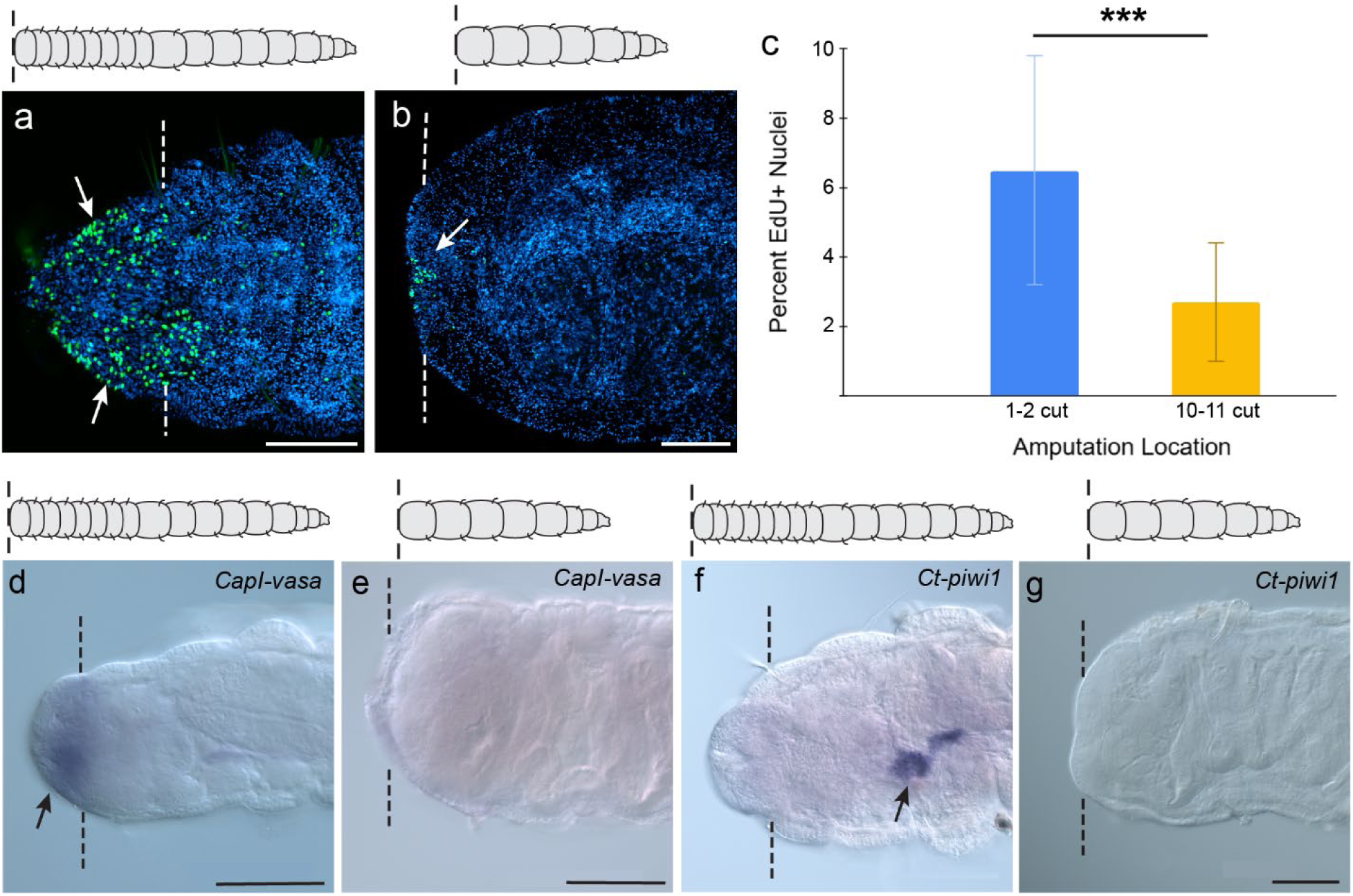
1-2 amputees show signs of blastema formation but 10-11 amputees do not. a-b) EdU incorporation (green) and nuclei stained with Hoechst 33342 (blue) in 3 dpa tail fragments cut between segments 1 and 2 (a) or 10 and 11 (b). White arrows point to areas of EdU incorporation. c) Quantification of EdU at 3 dpa in tail fragments cut between segments 1 and 2 or between segments 10 and 11. Sample sizes: 1-2 amputees: n=25, 10-11 amputees: n= 32. *** denotes statistical significance between groups with a p-value less than 0.05 using a Wilcoxon rank sum test. Error bars depict standard error. d-e) *CapI-vasa* expression is detected at the amputation site in 3 dpa 1-2 amputees (n= 27/32, black arrow, d), but not in 10-11 amputees (n= 27/29, e). f) *Ct-piwi1* expression is detected in the MPC cluster of 3 dpa 1-2 amputees, but not at the amputation site (n= 25/31). Black arrow points to *Ct-piwi1* expression in the MPC cluster. g) *Ct-piwi1* expression is not detected at the amputation site of 10-11 amputees (n= 20/29). In all images, anterior is to the left and view is ventral, except panel d, which is lateral. Dotted lines indicate amputation site. Scale bars, 100 µm.

Another characteristic of a regeneration blastema in *C. teleta* is expression of stem cell markers, such as *CapI-vasa*. At 3 dpa, 1-2 amputees display *CapI-vasa* expression at the amputation site (n= 27/32, Fig. 2d), whereas *CapI-vasa* expression is undetectable in 10-11 amputees (n= 27/29, Fig. 2e), in accordance with previous results (Kunselman & Seaver, 2025). In uncut animals, *CapI-vasa* expression is found in the posterior growth zone (pgz), in the MPC cluster in segment 5, and in coelomic cells posterior to segment 5 (de Jong & Seaver, 2018; Dill & Seaver, 2008). Since *CapI-vasa* is not expressed at the border of segment 1 and 2 in uncut animals, the *CapI-vasa* expression near the amputation site at 3 dpa likely arises *de novo* as a response to amputation in 1-2 amputees. Interestingly, *Ct-piwi1*, another stem cell marker, is not detectable at the amputation site of 3 dpa 1-2 amputees (n= 25/31) or 10-11 amputees (n= 29/29), which marks the first known instance in *C. teleta* where *Ct-piwi1* is not co-expressed with *CapI-vasa* in juveniles (Fig. 2f-g). *Ct-piwi1* and *CapI-vasa* are co-expressed in the MPC cluster and the pgz in intact juveniles, and in the posterior-facing blastema of regenerating head fragments.

Next, signs of cell differentiation were assessed at 7 dpa. Pax is a transcription factor and a marker of differentiated neurons (Webster et al., 2021). Two domains of Pax labeling are observed in 1-2 amputees: ventral and distal to the severed VNC but on the proximal side of the regenerating tissue (n= 18/26, Fig. 3a, arrow), and dorsal to the VNC (n= 7/26, Fig. S2, arrows). Dorsal Pax expression is colocalized with FMRFamide axon extensions that extend over the face of the gut proximal to the amputation site and therefore is likely attributed to the pre-existing stomatogastric ganglia rather than *de novo* dorsal expression (Fig. S2, arrows; Meyer et al., 2015). In contrast, 10-11 amputees only exhibit Pax labeling in nuclei of the pre-existing ganglia of the VNC at 7 dpa (n= 25/27, Fig. 3b). In uncut animals, Pax is expressed in the brain and ganglia of the VNC, and in the two stomatogastric ganglia that are bilaterally positioned at the dorsal posterior margin of the pharynx (n= 15/15, Fig. 3c-d; Meyer et al., 2015). Therefore, the ventral domain of Pax expression distal to the amputation site in 7 dpa 1-2 amputees arises in response to amputation, but the dorsal domain was likely present before amputation. *Ct-neuroD* is a gene that is expressed in neural cells that have exited the cell cycle (Sur et al., 2017), and it is expressed near the amputation site of 1-2 amputees at 7 dpa (n= 17/28). Usually *Ct-neuroD* expression is detected ventrally on the proximal side of the blastema (n= 10/28, Fig. 3e) or in several cells in the ganglion closest to the amputation site (n= 5/28). However, *Ct-neuroD* expression is undetectable near the amputation site of most 10-11 amputees (n= 30/32, Fig. 3f). Expression is limited to a few cells in the ganglion proximal to the amputation site in the two cases of 10-11 amputees that showed visible *Ct-neuroD* expression. In uncut animals, *Ct-neuroD* is expressed in the pgz (n= 37/38), in some cells of the enteric nervous system posterior to the pharynx (n= 27/38), and in a few cells in the brain (n= 11/38, Fig. S3), but not at the border of segments 1 and 2 or 10 and 11 (Fig. 3g-h). Thus, the *Ct-neuroD* expression observed near the amputation site in 1-2 amputees most likely arises after amputation. The homeobox gene *orthodenticle* (*otx*) plays a conserved role in head patterning and neurogenesis (Bely & Wray, 2001; Finkelstein & Perrimon, 1990; Simeone et al., 1993). This is the case in *C. teleta* because *Ct-otx* is expressed in the anterior ectoderm associated with the brain anlagen in an early-stage larva, and then in the brain, foregut, VNC, and pgz as a late-stage larva (Bely & Wray, 2001; Boyle et al., 2014). Following amputation between segments 1 and 2, *Ct-otx* is expressed at the proximal side of the blastema (n= 8/22) or dorsal to the gut protrusion (n= 4/22, Fig. 3i). In contrast, *Ct-otx* expression is undetectable in 10-11 amputees at the cut site (n= 26/27, Fig. 3j), with the exception of one case with faint expression in the mesoderm at the amputation site. In uncut juveniles, *Ct-otx* is faintly expressed in the brain (n= 30/37), pgz mesoderm (n= 25/32), and foregut (n=37/37), but its expression at the boundary of segments 1 and 2 and segments 10 and 11 is weak and undetectable, respectively (n= 37/37, Fig. 3k-l). When *C. teleta* head fragments regenerate a tail, *Ct-otx* is also strongly expressed bilaterally in the mesoderm of the blastema (n= 11/23) or broadly across the distal edge of the blastema in the mesoderm at 3 dpa (n= 10/23, Fig. S4).

**Figure 3.**
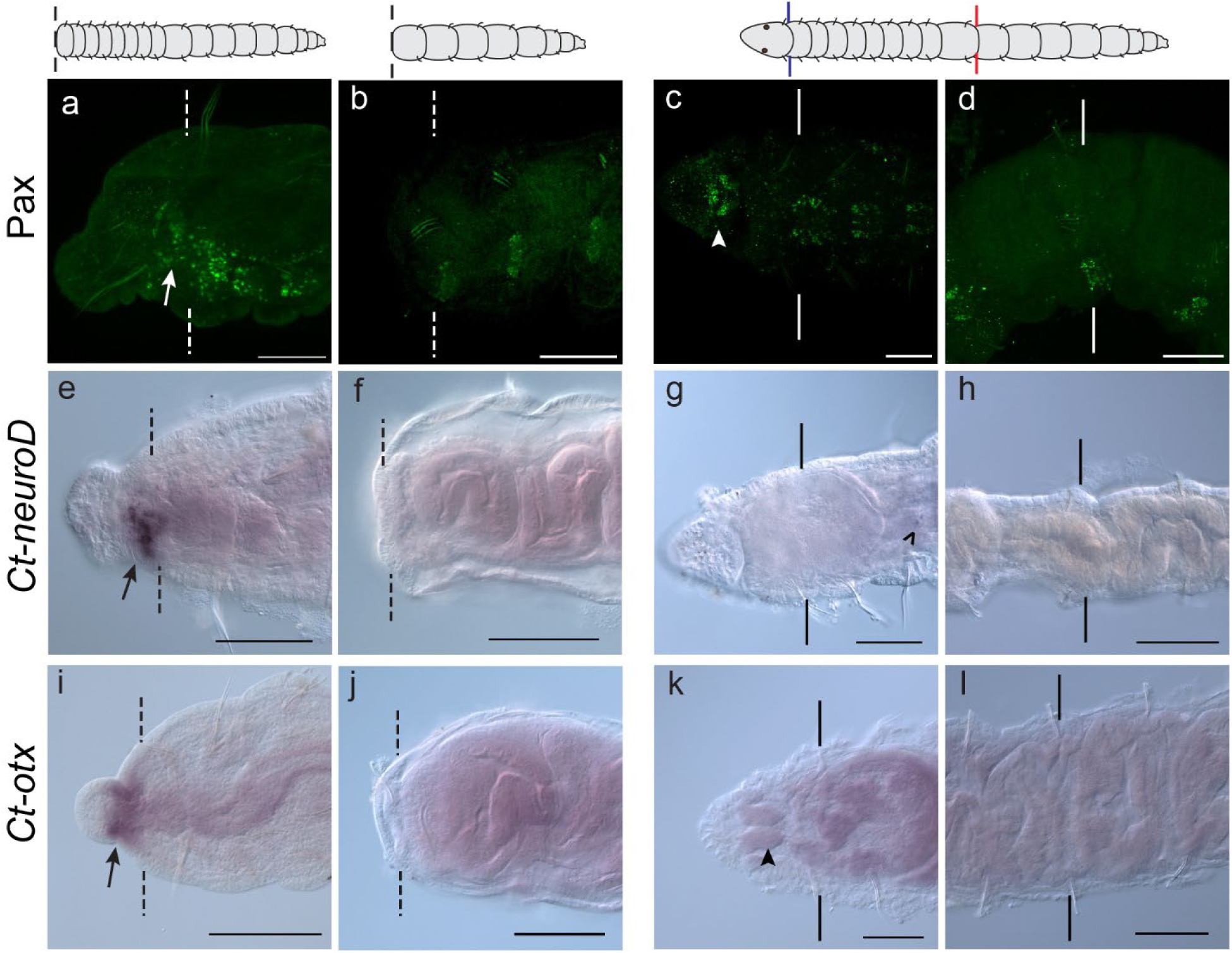
1-2 amputees show signs of neural differentiation but 10-11 amputees do not. a-d) Anti-Pax antibody (green) labels nuclei of cells specified to become neural cells at the amputation site of 1-2 amputees (a) and 10-11 amputees (b), at the border between segments 1 and 2 of uncut animals (c), and at the border between segments 10 and 11 in uncut animals (d). White arrow points to anti-Pax+ cells distal to the severed VNC. White arrowhead points to the brain. e-h) *Ct-neuroD* expression at the amputation site 1-2 amputees (e) and 10-11 amputees (f) at 7 dpa, and at the border between segments 1 and 2 of intact animals (g), and at the border between segments 10 and 11 in uncut animals (h). i-l) *Ct-otx* is expressed at the amputation site 1-2 amputees (i) and 10-11 amputees (j) at 7 dpa, at the border between segments 1 and 2 of uncut animals (k), and at the border between segments 10 and 11 in uncut animals (l). Black arrows indicate localized gene expression at the amputation site. Open arrowhead indicates *Ct-neuroD+* cells in the enteric nervous system posterior to the pharynx. Solid black arrowhead points to the brain. Dotted lines indicate amputation site. Solid lines indicate a segment boundary. In all images, anterior is to the left. In panels a, b, d, e, and j view is lateral. In panel g view is ventral. In remaining panels, view is dorsal. Scale bars, 100 µm.

To visualize the morphology of the nervous system at 7 dpa to see if 1-2 amputees regenerate differentiated structures typically found in the head, an acetylated tubulin antibody was used to label all nerves, and cross-reactive antibodies against serotonin and FMRFamide were used to visualize distinct neural subtypes (Meyer et al., 2015). At the boundary of segments 1 and 2 in intact animals, two thick circumesophageal connectives extend anteriorly from the first ganglion of the VNC around the mouth, where they connect to each lobe of the brain (open arrowheads, Fig. 4a-b; Meyer et al., 2015). By 7 dpa, acetylated tubulin labeling in 1-2 amputees reveals abundant and highly branching nerve extensions from the VNC that densely fill the regenerating tissue, rather than two thick connectives reminiscent of the circumesophageal connectives (Fig. 4c). Occasionally, sensory structures that look like tufts of cilia in the epidermis form at the distal end of the blastema in 1-2 amputees (solid arrowheads, n= 5/21, Fig. 4c). In uncut animals, these structures are densely packed at the anterior tip of the worm as well as in the pygidium (solid arrowheads, Fig. 4a-b). The peripheral nerves adjacent to the amputation site also project axons anteriorly towards the wound in 1-2 amputees. Branching of the peripheral nerves is not observed during posterior regeneration in *C. teleta*, rather, only the longitudinal nerves of the VNC project into the blastema (de Jong & Seaver, 2016). In accordance with previous results, there are some neurite extensions from the VNC over the wound face in 10-11 amputees (n= 23/27), but this was limited to about four to five very fine neurites when present, and contrasts with the copious, meandering projections at the wound site of 1-2 amputees (Fig. 4d). In uncut animals, there are serotonin-positive cell bodies in the brain and they project axons along the circumesophageal connectives and the connectives of the VNC, where additional serotonin-positive cell bodies are present in the ganglia (Fig. 4e-f; Meyer et al., 2015). In 1-2 amputees, serotonin-positive neurites project dorsally from the VNC (n= 20/21, Fig. 4g, arrow). In 10-11 amputees, very fine serotonin-positive neurite extensions were visible across the wound face in some cases (n= 12/27), but in other cases, no serotonin-positive extensions from the VNC were visible (n= 15/27, Fig. 4h). FMRFamide labels cell bodies in the ganglia of the VNC, axon projections in the circumesophageal connectives and VNC, as well as a pair of anterior enteric nerves (aENs) that project along the dorsal-lateral side of the dorsal pad of the pharynx and terminate at the stomatogastric ganglia (Fig. 4i-j; Meyer et al., 2015). In 1-2 amputees, FMRFamide-positive neurites appear to extend from the VNC, sometimes projecting laterally around the gut protrusion (n= 7/26) or anteriorly into the gut protrusion or the blastema (n= 9/26), and other times projecting dorsally and posteriorly towards the stomatogastric ganglia (n= 11/26, Fig. 4k, arrow). In 10-11 amputees, FMRFamide-positive neurite outgrowths from the VNC are not detected in the majority of cases (n= 20/27, Fig. 4l). Taken together, pre-existing peripheral nerves, serotonin-positive neurons, and FMRFamide-positive neurons send out more projections in 1-2 amputees than 10-11 amputees, but the projections do not appear to be patterned like the nervous system in the head of uncut animals.

**Figure 4.**
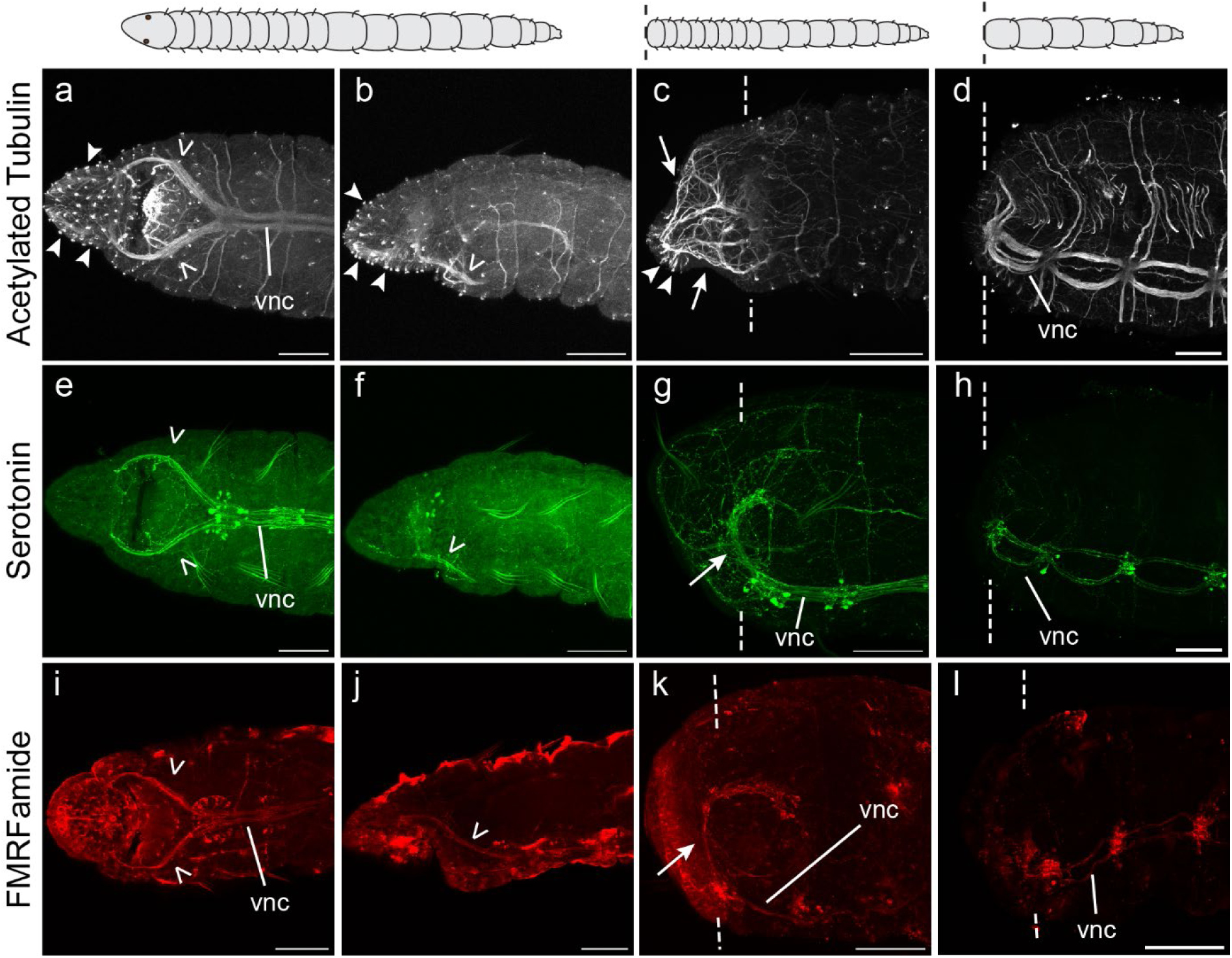
Anterior nervous system anatomy and nervous system response to amputation at two distinct amputation sites. a-d) Anti-acetylated tubulin (white) labels the nervous system of intact animals (a-b), tail fragments cut between segments 1 and 2 (c), and tail fragments cut between segments 10 and 11 (d). Solid arrowheads point to ciliated sensory structures. Open arrowheads point to circumesophageal connectives. Arrows point to neurites in regenerated tissue. e-h) Anti-serotonin (green) labels a subset of neurons of uncut animals (e-f), tail fragments cut between segments 1 and 2 (g), and tail fragments cut between segments 10 and 11 (h). Open arrowheads point to circumesophageal connectives. Arrow points to dorsal neurite extensions from the VNC. i-l) Anti-FMRFamide (red) labels a subset of neurons in intact animals (i-j), tail fragments cut between segments 1 and 2 (k), and tail fragments cut between segments 10 and 11 (l). Open arrowheads point to circumesophageal connectives. Arrow points to dorsal neurite extensions from the VNC. Dotted vertical lines indicate amputation site. In all images, anterior is to the left. In first column, view is ventral. In all other panels, view is lateral. Scale bars, 100 µm. Abbreviations: vnc, ventral nerve cord.

In total, the higher number of proliferating cells, presence of stem-cell marker expression, and expression of neural differentiation markers proximal to the amputation site of 1-2 amputees compared to 10-11 amputees demonstrate that 1-2 amputees have greater regeneration potential than 10-11 amputees. However, not all missing structures in the head are replaced by 7 dpa; therefore, regeneration does not go to completion.

### Vasa-positive Coelomic Cells Are Rarely Found Anterior to the MPC Cluster

When hypothesizing reasons why regeneration potential is higher in 1-2 amputees compared to 10-11 amputees, the putative stem cell cluster (the MPC cluster) located in segment 5 of *C. teleta* seems like a logical cause. Previous experimental manipulations suggest that presence of the MPC cluster contributes to posterior regeneration in *C. teleta,* most likely via cells migrating through the coelomic cavity from the MPC cluster to the cut site (de Jong & Seaver, 2018). Since the MPC cluster is located in segment 5, 1-2 amputees retain the MPC cluster following amputation, while the MPC cluster is removed in 10-11 amputees as a result of amputation. However, individual *CapI-vasa*-positive cells are rarely detected anterior to the MPC cluster in the coelomic cavity of juveniles amputated posterior to segment 10 over the course of regeneration (de Jong & Seaver, 2018). Therefore, to determine whether a transverse cut made anterior to the cluster can stimulate Vasa-positive cells (putatively from the MPC cluster) to appear in the body between the MPC cluster and the amputation site, we used an antibody against Vasa to monitor presence of these cells in 1-2 amputees following amputation.

In uncut animals, *CapI-vasa* is detectable in the MPC cluster, cells in the coelomic cavity, and the pgz (de Jong & Seaver, 2018). During posterior regeneration, the Vasa antibody labels the MPC cluster, cells in the coelomic cavity, and a broad domain in the blastema (Fig. 5a). *CapI-vasa*-positive coelomic cells posterior to the MPC cluster are present at a higher frequency in regenerating head fragments at 2 and 3 dpa than in intact juveniles (de Jong & Seaver, 2018). After cutting anterior to the MPC cluster between segments 1 and 2, nine or fewer Vasa-positive coelomic cells are detected between the MPC cluster and the edge of the wound 3 dpa (n= 22/22, Fig. 5b). On average, two Vasa-positive cells are detected anterior to the MPC cluster and in 9/22 cases, no Vasa-positive cells were observed anterior to the MPC cluster. These few cells can be found randomly dispersed across the entire span of space between the distal wound edge and the MPC cluster. Faint Vasa expression at the cut site, reminiscent of broad Vasa expression in the blastema during posterior regeneration, is visible, as well as numerous Vasa-positive coelomic cells posterior to the MPC cluster (Fig. 5b). Vasa-positive coelomic cells are also observed in 3 dpa 10-11 amputees along the length of the tail, although these may have been present before amputation (n= 32/32, Fig. 5c). These data suggest that Vasa-positive coelomic cells do not make a substantial contribution to the increased regeneration ability of 1-2 amputees. This observation marks an additional difference between anterior-facing and posterior-facing blastemas.

**Figure 5.**
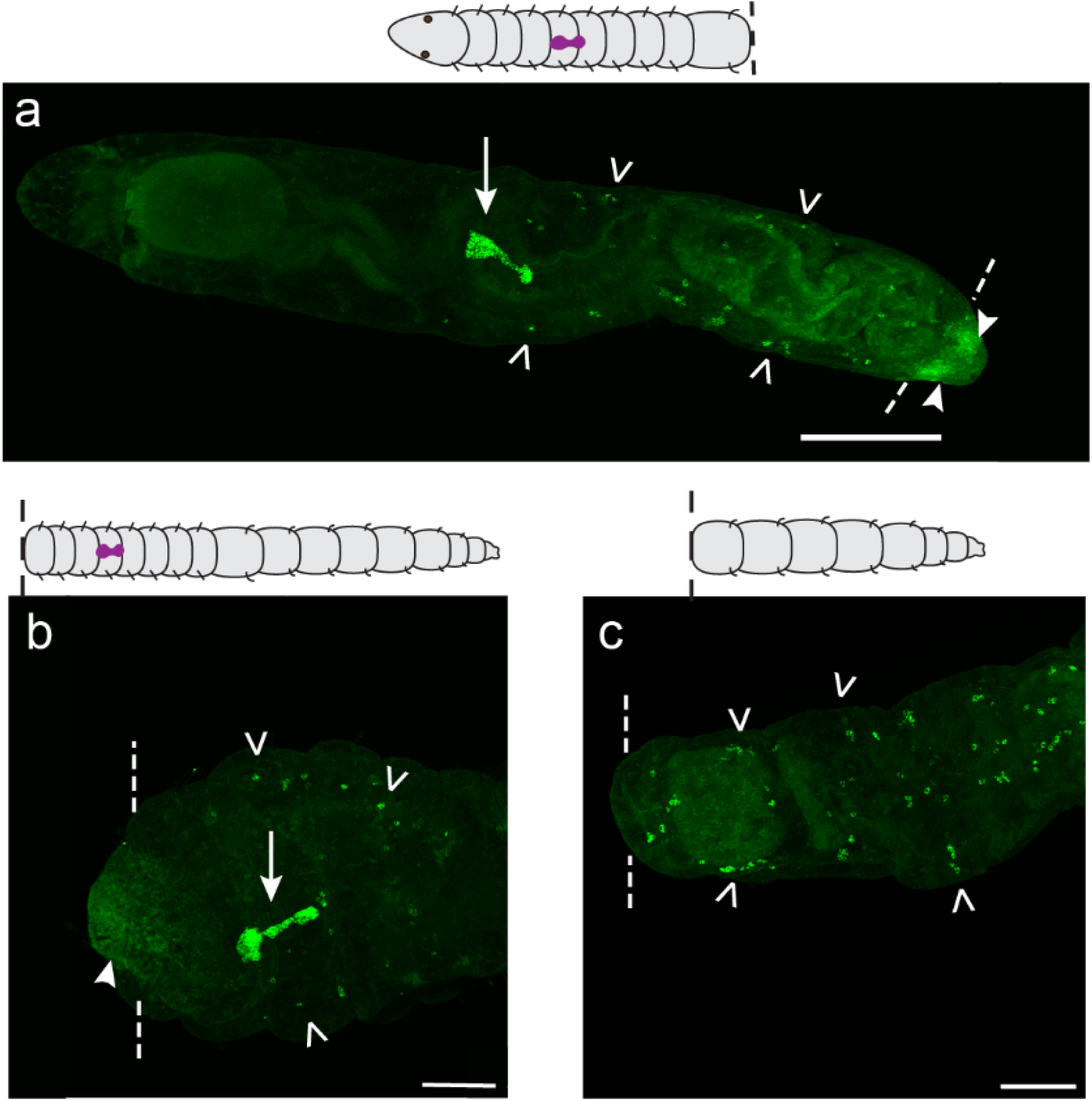
Vasa+ cells in the coelom are rarely detected anterior to the MPC cluster. a) Vasa expression in 3 dpa head fragments cut between segments 10 and 11. b) Vasa expression in tail fragments cut between segments 1 and 2. c) Vasa expression in tail fragments cut between segments 10 and 11. Arrows point to the MPC cluster. Open arrowheads point to cells in the coelomic cavity. Solid arrowheads point to Vasa expression in the blastema. In all images, anterior is to the left and view is ventral. Dotted lines indicate amputation site. Scale bars, 200 µm (a), 100 µm (b-c).

Although an anterior-facing wound does not attract or stimulate production of Vasa-positive coelomic cells, we wondered if these cells are influenced by Wnt/β-catenin signaling since components of this signaling pathway are highly upregulated in posterior-facing blastemas of *C. teleta* (Kunselman & Seaver, 2025). To test this hypothesis, 1-2 amputees were treated with a Wnt/β-catenin signaling agonist, CHIR-98014 (CHIR) for three consecutive days following amputation and then fixed. All DMSO controls and CHIR-treated 1-2 amputees have fewer than ten Vasa-positive coelomic cells between the wound edge and the MPC cluster (Fig. 6). DMSO controls have an average of three Vasa-positive cells anterior to the MPC cluster (n=33), and CHIR-treated amputees have an average of 2.5 cells anterior to the MPC cluster (n=35). Therefore, increased Wnt/β-catenin signaling does not affect the number of Vasa-positive coelomic cells anterior to the MPC cluster.

**Figure 6.**
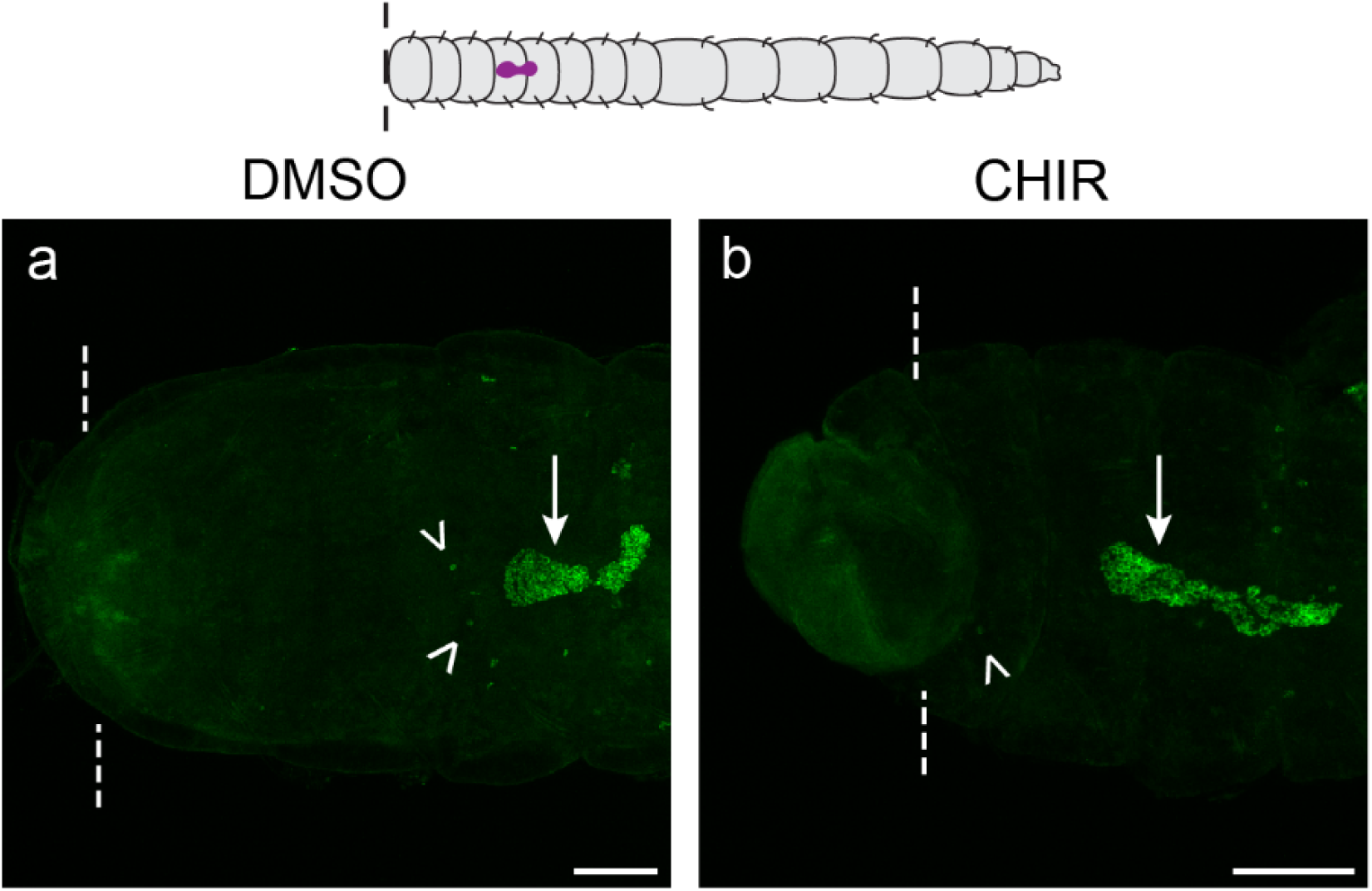
1-2 amputees treated with CHIR rarely have Vasa+ coelomic cells anterior to the MPC cluster. a) Anti-Vasa expression in 3 dpa DMSO-treated control tail fragments cut between segments 1 and 2. b) Anti-Vasa expression in CHIR-treated tail fragments cut between segments 1 and 2. Arrows point to cells in the MPC cluster. Open arrowheads point to cells in the coelomic cavity anterior to the MPC cluster. All images are in ventral view with anterior to the left. Dotted lines indicate amputation site. Scale bars, 100 µm.

### Cutting Proximal to the MPC Cluster Does Not Lead to Complete Regeneration

We next asked whether regeneration of tail fragments could go to completion if Vasa-positive coelomic cells were present at the amputation site. To do this, an amputation was made between segments 4 and 5, immediately adjacent to and along the anterior edge of the MPC cluster. As a result of this amputation, the cells are located at the wound site. Tail fragments that were amputated between segments 4 and 5 will be referred to as 4-5 amputees henceforth. The MPC cluster is present in almost all animals amputated between segments 4 and 5 (n= 13/15), demonstrating that amputation at this position does not inadvertently remove the MPC cluster (Fig. 7a). Not surprisingly, in 4-5 amputees, the MPC cluster persists until 7 dpa, although there is scarcely any new tissue regenerated and they appear comparable to specimens fixed immediately following amputation (Fig. 7b-h). The 4-5 amputees wound heal (Fig. 7f) and exhibit some cell division (Fig. 7g) and axon extension (Fig. 7h) at the amputation site, but the amount of regenerated tissue distal to the amputation site is inconsequential. Therefore, the presence of the MPC cluster at the wound does not rescue regeneration.

**Figure 7.**
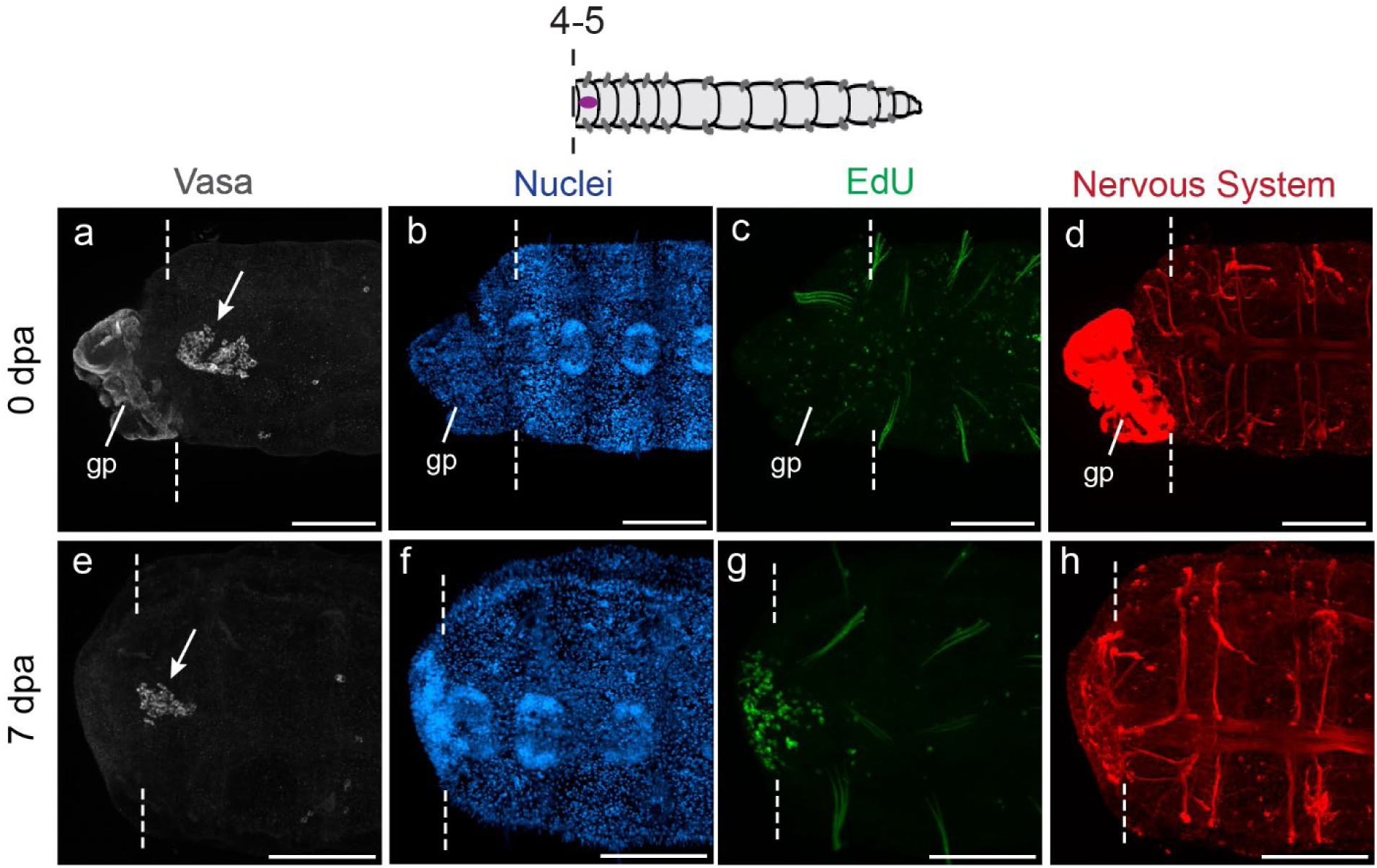
Amputating at the MPC cluster does not rescue regeneration. a-d) Tail fragment cut between segments 4 and 5 and fixed within one hour of amputation. e-h) Tail fragment cut between segments 4 and 5 and fixed 7 dpa. Anti-vasa labeling in white, nuclei stained with Hoechst 33342 in blue, EdU incorporation in green, and anti-acetylated tubulin in red. (a-d) and (e-h) are images of the same individual. Arrows point to the MPC cluster. Dotted lines indicate amputation site. All images are in ventral view with anterior to the left. Scale bars, 100 µm. Abbreviations: gp, gut protrusion.

### Activating Wnt/β-catenin Signaling Does Not Lead to Further Differentiation

In a previous study, we showed that experimentally activating Wnt/β-catenin in tail fragments amputated between segment 10 and 11 leads to completion of early, but not late, stages of posterior regeneration (Kunselman & Seaver, 2025). Since we demonstrate in this study that 1-2 amputees have more inherent regeneration potential than 10-11 amputees, we hypothesized that activation of Wnt/β-catenin signaling in 1-2 amputees would lead to complete posterior regeneration. 1-2 amputees were treated with CHIR for 0-3 dpa, then washed out of the drug and raised for an additional four days and fixed at 7 dpa. The 0-3 dpa treatment interval was chosen because in 10-11 amputees, treatment from 0-3 dpa was sufficient to induce the same phenotype as 10-11 amputees treated with the Wnt/β-catenin agonist for 7 days continuously (Kunselman & Seaver, 2025). Most 10-11 amputees treated with CHIR 0-3 dpa rarely show signs of later steps of regeneration such as formation of new ganglia (n= 3/31), a subterminal band of EdU-positive cells (n= 3/31), or new peripheral nerves (n= 4/31, Fig. 8a-c). 1-2 amputees treated with CHIR from 0-3 dpa form tissue distal to the cut site with EdU incorporation at the wound site (n= 40/40, arrow), neurite extensions (n= 38/40), and occasionally tuft-like sensory structures typically found on the terminal ends of intact animals (n= 17/40, arrowheads, Fig. 8d-f). However, these animals rarely form a bilateral, subterminal band of EdU-positive cells typical of a pgz (n= 3/40), or new ganglia (n= 0/40) and peripheral nerves (n= 1/40), all of which are indicators of newly formed segments. A longer, continuous CHIR incubation for 7 dpa also does not lead to formation of ganglia, peripheral nerves, or a pgz in 1-2 amputees (n= 21/21, Fig. 8g-i). This suggests that tissue with a higher inherent regeneration potential is not guaranteed to produce a successful regeneration outcome upon activation of Wnt/β-catenin signaling.

**Figure 8.**
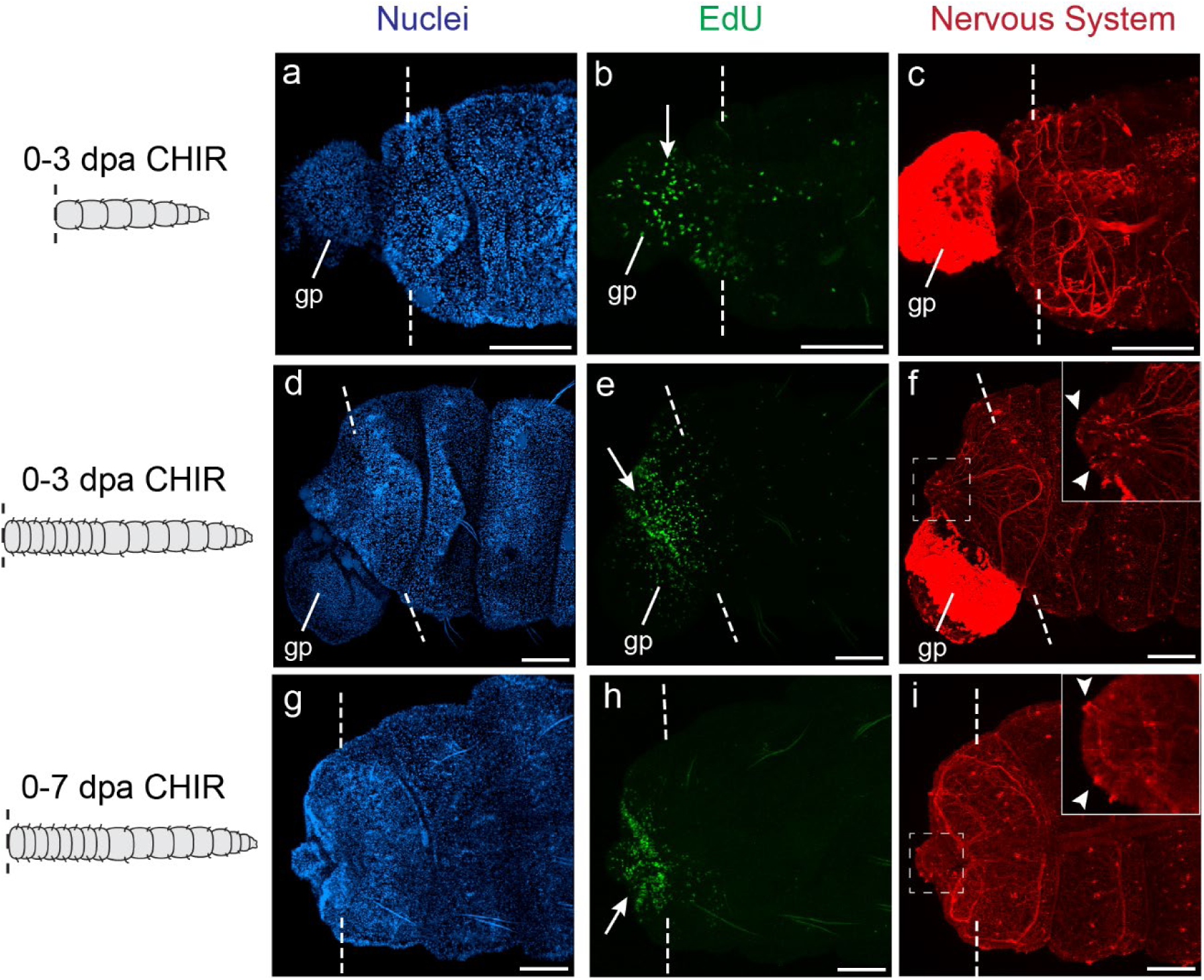
Activation of Wnt/β-catenin signaling does not increase regenerative potential of 1-2 amputees. a-c) Tail fragment cut between segments 10 and 11, treated with CHIR 0-3 dpa, and fixed 7 dpa. n= 31. d-f) Tail fragment cut between segments 1 and 2, treated with CHIR 0-3 dpa, and fixed 7 dpa. n= 40. g-i) Tail fragment cut between segments 1 and 2, treated with CHIR 0-7 dpa, and fixed 7 dpa. n=21. Nuclei stained with Hoechst 33342 in blue, EdU incorporation in green, and anti-acetylated tubulin in red. (a-c), (d-f), and (g-i) are images from the same individual. Arrows point to EdU localized to amputation site. Arrowheads point to regenerated sensory tufts. Dotted lines indicate amputation site. Dotted box indicates area shown in inset. In all images, anterior is to the left. In top and middle row, view is lateral. In bottom row, view is ventral. Scale bars, 100 µm.

Surprisingly, two 10-11 amputees treated with CHIR 0-3 dpa from two independent experiments exhibited three major characteristics of complete posterior regeneration: formation of new ganglia, a subterminal band of EdU-positive cells typical of a pgz, and new peripheral nerves (n= 2/31, Fig. 9). Therefore, activation of Wnt/β-catenin signaling in 10-11 amputees is sufficient to induce complete posterior regeneration, although the event is relatively rare (n= 2/31).

**Figure 9.**
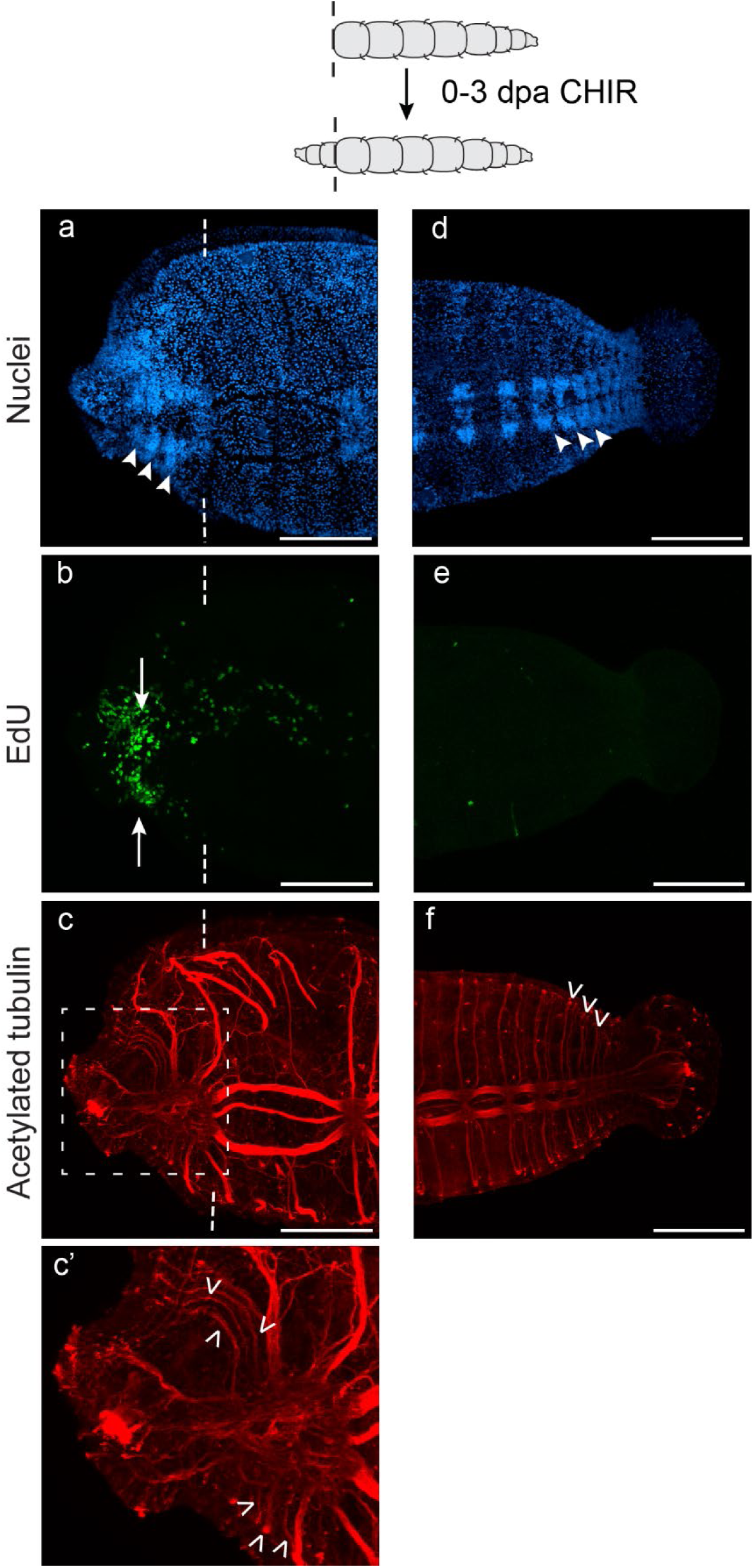
Complete posterior regeneration after Wnt/β-catenin activation in a tail fragment cut between segments 10 and 11. Tail fragments were treated with CHIR 0-3 dpa and raised to 7 dpa. a-c) The anterior-facing end of the 10-11 cut. a) Nuclei stained blue with Hoechst 33342. Arrowheads point to regenerated ganglia. b) EdU incorporation in green. Arrows point to bilateral, subterminal domains of EdU incorporation near the amputation site. c) Anti-acetylated tubulin in red labels the nervous system. c’) Inset of area inside the dotted box in panel c. Open arrowheads point to regenerated peripheral nerves. d-f) Posterior-facing end of the 10-11 cut. d) Nuclei stained blue with Hoechst 33342. Arrowheads point to ganglia in the posterior of the body fragment. e) EdU incorporation is in green. f) Anti-acetylated tubulin in red labels the nervous system. Open arrowheads point to segmentally repeated peripheral nerves. Dotted lines indicate amputation site. All images are in ventral view and anterior is to the left. Scale bars, 100 µm.

## Discussion

### Regeneration Potential Varies Along the AP axis of *C. teleta*

Gradations of regeneration potential along the AP axis are well documented in many species of annelids, but this study is the first to report the regeneration ability of *C. teleta* tail fragments at different amputation positions. Our descriptions of cell division patterns and stem cell gene expression suggest that 1-2 amputees form an anterior-facing blastema, while 10-11 amputees only wound heal and show limited neurite extension. Furthermore, there is evidence of neural differentiation in the regenerating tissue of 1-2 amputees. Our ability to score for subtle neural phenotypes like the VNC neurite extensions to the stomatogastric ganglia is made possible by usage of cross-reacting antibodies that label neural subtypes, which in this case gives a clearer picture of the nervous system’s response to amputation than does labeling the entire nervous system with acetylated tubulin. Overall, 1-2 amputees proceed to later stages of regeneration than 10-11 amputees, suggesting that the anterior tissue abutting the amputation plane has greater regeneration potential than the abdominal tissue where 10-11 amputees are severed.

Variation in regeneration potential across the AP axis is common in many annelids (Berrill, 1928, 1952; Hyman, 1940; Myohara, 2012; Planques et al., 2019; Spieß et al., 2024). Through documentation of regeneration characteristics in many annelid species, historical literature confirms that anterior regeneration ability generally deteriorates or weakens as amputations are made further posterior (Hyman, 1940). For example, *C. variopedatus* anterior regeneration potential abruptly terminates posterior to segment 14 (Berrill, 1928). In lumbricid earthworms, fewer anterior segments are regenerated as amputations are made further posterior, until no new segments form anteriorly (Hyman, 1940). While we do not have evidence that the anterior-facing blastema in 1-2 amputees has an anterior identity, we believe it is likely, for reasons described later in this discussion. If the blastema of 1-2 amputees does have anterior identity, our findings suggest head regeneration in *C. teleta*, although incomplete, proceeds to later stages of regeneration at more anterior amputation positions. In this way, *C. teleta* is similar to other annelids that have stronger anterior regeneration ability at more anterior amputation sites.

The mechanisms explaining differences in regeneration potential along the AP axis in annelids are still largely unknown. One hypothesis put forth by the literature is that some subregions of the gut are susceptible to extrude from the wound and negatively impact regeneration. The proventricle region of the gut in *S. malaquini* is proposed to inhibit anterior and posterior regeneration because its protrusion after amputation hinders wound healing, leading to high mortality (Spieß et al., 2024). Posterior regeneration is inhibited as amputations are made within one segment of the pharynx in *P. dumerilii* because the protrusion of the pharynx through the wound leads to high mortality (Planques et al., 2019). In contrast, our amputations between segments 1 and 2 are adjacent to the pharynx and have more regeneration potential than 10-11 amputees amputated in the midgut region. We see protrusion of the gut to seal the wound in both 1-2 and 10-11 amputees. Gut protrusions are uncommon in *C. teleta* head fragments, which possess robust posterior regeneration ability (Kunselman & Seaver, 2025). However, gut protrusion and blastema formation are not mutually exclusive in *C. teleta* and their co-occurrence has been documented (Kunselman & Seaver, 2025). Although wound healing is important for regeneration to proceed and the gut certainly plays a role in this process, improper wound healing does not explain why 10-11 amputees fail to regenerate an anterior-facing blastema, since they wound heal and can survive up to a week following amputation.

Another tissue proposed to regulate annelid regeneration abilities is the central nervous system. In *P. dumerilii*, the presence of the brain is necessary for posterior regeneration to occur, and it is hypothesized that the brain produces a growth-promoting hormone present in a graded manner along the AP axis, which explains why posterior regeneration occurs at a faster rate when amputations are made progressively more anterior (Planques et al., 2019). In the earthworm *Eisenia foetida*, implanting or deflecting the VNC towards the body wall forms ectopic segments with a brain, and the inductive power of the VNC is strongest at the anterior end and gradually becomes weaker toward the posterior (Berrill, 1952; Okada & Kawakami, 1943). There are some differences in VNC structure between thoracic and abdominal segments in *C. teleta* (e.g. size of ganglia, distance between left and right connectives), and *Hox* genes are expressed regionally along the VNC with discrete anterior and posterior expression boundaries (de Jong & Seaver, 2016; Meyer et al., 2015). This variation in morphology and gene expression along the length of the VNC may impact different regeneration responses depending on where the VNC is severed. Although we certainly see differences in the pattern of axon extensions between 1-2 and 10-11 amputees, it is unclear how or if the VNC influences the formation of the anterior-facing blastema or the limited neural differentiation proximal to the amputation site.

An exciting frontier to consider as an underlying mechanism of variable regeneration ability along the AP axis in annelids is chromatin remodeling. After injury, many epigenetic changes take place in cells to activate transcriptional programs for tissue regeneration (Rodriguez & Kang, 2020; Rouhana & Tasaki, 2016). Regeneration-specific enhancers have been identified and, in some cases, validated in a variety of organisms (Cazet et al., 2021; Gehrke et al., 2019; Jimenez et al., 2022; Loubet-Senear & Srivastava, 2024; Rodriguez & Kang, 2020). ATAC-seq is a reliable method to identify genome-wide changes in accessible chromatin (Buenrostro et al., 2013), but to date the only ATAC-seq study performed in annelids focused on development and not regeneration (Martín-Zamora et al., 2023). ATAC-seq of 1-2 amputees and 10-11 amputees in *C. teleta* could reveal differences in chromatin remodeling around promoters or enhancers driving variation in regeneration potential between the two amputation locations.

Another way to explore the temporal and spatial dynamics of chromatin remodeling is by labeling regenerating tissue with antibodies against K9 dimethylated histone H3, acetylated histone H3, or acetylated histone H3 lysine 9 during regeneration (Niwa et al., 2013; Suzuki et al., 2016). The former marker is associated with transcriptionally inactive, differentiated cells, while the latter markers are associated with actively transcribed chromatin (Niwa et al., 2013; Suzuki et al., 2016). A study in *Xenopus laevis* developed transgenic larvae that ubiquitously express a fluorescent antibody that binds to acetylated histone H3 lysine 9, enabling visualization of the dynamics of active chromatin *in vivo* during tail regeneration (Suzuki et al., 2016). Performing a similar experiment in *C. teleta* at different amputation sites could reveal interesting patterns in acetylation dynamics in cells along the AP axis.

### Anterior-facing Blastemas Largely Lack Coelomic Vasa-positive Cells

Resident adult stem cells are sources of regenerated tissue in many highly regenerative organisms such as the cnidarian *Hydractinia symbiolongicarpus*, the acoel *H. miamia*, the planarians *S. mediterranea* and *Dugensia japonica*, and the tapeworm *Hymenolepis diminuta* (Gehrke & Srivastava, 2016; Guedelhoefer & Alvarado, 2012; Orii et al., 2005; Reddien & Sánchez Alvarado, 2004; Varley et al., 2023; Wenemoser & Reddien, 2010). We hypothesized that the increased regenerative potential of 1-2 amputees could be due to the migration of Vasa-positive stem cells from the MPC cluster located in segment 5 towards the wound. There is strong experimental evidence that the MPC cluster contributes to posterior regeneration in *C. teleta* (de Jong & Seaver, 2018). The MPC cluster is retained in tail fragments generated from cuts between segments 1 and 2, but is removed during amputation in 10-11 amputees. Additionally, there is a precedent for stem cell-based regeneration in annelids. Cells with stem cell-like characteristics were described as early as 1892 in the annelid *Lumbriculus*, and based on histological sections, these cells, termed neoblasts, were inferred to generate most of the new mesoderm (Randolph, 1892). Migration of cells with neoblast-like qualities towards amputation wounds has been recorded by time lapse imaging of the annelid *P. leidyi*, although the identity of migratory cells and their contribution to the regenerated tissue is uncertain (Zattara et al., 2016). Neoblasts have also been reported to contribute to the regeneration blastema in the annelid *Enchytraeus japonensis* (Myohara, 2012; Tadokoro et al., 2006). In the historical literature, Berrill (1952) reviews descriptions of neoblasts migrating posteriorly towards a wound in the annelids *Aricia*, *Chaetopterus*, *Diopatra*, *Tubifex*, and *Nais*. Interestingly, however, he points out that neoblasts are never described as migrating anteriorly towards an anterior wound in any of these species (Berrill, 1952).

When investigating whether Vasa-positive coelomic cells in *C. teleta* contribute to the blastema of tail fragments cut between segments 1 and 2, we detect very few Vasa-positive coelomic cells between the amputation plane and the MPC cluster. This finding suggests that these cells do not significantly contribute to the generation of the anterior-facing blastema, although we cannot rule out the possibility that a few Vasa-positive coelomic cells may divide rapidly to generate the cells of the regenerating tissue. We feel this is unlikely given the vastly greater numbers of EdU-positive cells at the wound site compared to Vasa-positive cells (compare Fig. 2a to Fig. 5b). Furthermore, amputating immediately adjacent to the MPC cluster does not induce tissue regeneration distal to the amputation plane. Thus, although we cannot rule out transdifferentiation or contribution of stem cells that do not express *CapI-vasa*, we propose that anterior-facing blastemas are generated via dedifferentiation of cells proximal to the wound.

Additional experiments can clarify if transdifferentiation or other populations of stem cells serve as the source of cells during *C. teleta* regeneration. For example, treating *C. teleta* tail fragments with a cell-cycle inhibitor such as hydroxyurea and assessing whether neural specification occurs in the absence of cell division would provide evidence of transdifferentiation (Planques et al., 2019; Zheng et al., 2022). An EdU pulse in uncut worms followed by amputation, a chase of a few days, and then a BrdU pulse prior to fixation could demonstrate what proportion of the regenerate contains EdU-positive cells originating from the pre-existing tissue (de Jong & Seaver, 2018; Dirks et al., 2012; Planques et al., 2019; Ramon-Mateu et al., 2019). Irradiation of *C. teleta* prior to amputation to deplete resident stem cells may significantly inhibit regeneration if stem cells play an essential role in tissue regeneration (Gehrke & Srivastava, 2016; Guedelhoefer & Alvarado, 2012; Rozario et al., 2019). Development of transgenics to label stem cells or cells derived from different germ layers could reveal how different populations of cells respond to amputation, as well as what cell types they can produce (Ricci & Srivastava, 2021; Tanaka & Reddien, 2011; Zheng et al., 2022). In summary, there are many feasible methods available to explore the source of regenerated tissue in *C. teleta*.

### Activation of Wnt/β-catenin Signaling Has Little Effect on Vasa-positive Coelomic Cell Location

Stem cells may have to travel great distances towards a wound to supply a source of cells to a regeneration blastema, and therefore guidance cues are crucial for these cells to reach their target. In *C. teleta*, a transverse cut anterior to the MPC cluster results in few Vasa-positive cells positioned between the MPC cluster and the wound, suggesting that putative migratory Vasa-positive cells are not attracted to signals from an amputation wound located anterior of the MPC cluster. This result is surprising since Vasa-positive coelomic cells appear to migrate posteriorly from the MPC cluster during posterior regeneration (de Jong & Seaver, 2018). In the planarian *S. mediterranea*, tissue transplantation and partial irradiation experiments demonstrate that stem cells are mobilized upon tissue injury and migrate towards both anterior and posterior amputation sites (Guedelhoefer IV. & Alvarado, 2012). During homeostasis, the stem cells of *S. mediterranea* are relatively stationary (Guedelhoefer IV. & Alvarado, 2012). In mammals, mesenchymal stromal cells are multipotent adult progenitor cells that migrate to sites of injury using chemotactic signals released from damaged tissue such as platelet-derived growth factor-AB, insulin-like growth factor (IGF)-1, and stromal cell-derived factor-1 (SDF-1) (Ullah et al., 2019). Since a few cases of Vasa-positive cells were documented to be located anterior to the MPC cluster after amputation in *C. teleta*, these cells appear to have the capability to migrate anteriorly, if they are in fact migrating. Because Wnt/β-catenin signaling is upregulated during posterior regeneration and Vasa-positive coelomic cells are often found between the MPC cluster and the posterior wound edge, we hypothesized that Wnt/β-catenin signaling may exert an attractive influence on Vasa-positive coelomic cells. However, increasing Wnt/β-catenin signaling with CHIR treatment did not lead to increased instances of Vasa-positive cells detected between the MPC cluster and the wound edge of 1-2 amputees after 3 or 7 dpa. One limitation of this experiment is that CHIR was uniformly applied to tail fragments and not specifically applied at the wound; however, since we previously observed a localized response at the wound site following global application of CHIR (Kunselman & Seaver, 2025), we think that broad application of CHIR in this study did not greatly affect our results. In the literature, canonical Wnt signaling is recognized for its role in regulating cell proliferation and differentiation, so rather than guiding stem cells, it may help sustain their self-renewing properties and their undifferentiated character (Sonavane & Willert, 2023).

Aside from hematopoietic stem cells and mesenchymal stromal cells in mammals, little is known about the mechanisms that guide stem cells out of their niches towards injured tissue (Guedelhoefer IV. & Alvarado, 2012; Laird et al., 2008). Mechanisms that guide stem cells during development include G-protein coupled receptor (GPCR)-mediated guidance in *Drosophila melanogaster* primordial germ cells (Kunwar et al., 2003) and Eph receptor and ephrin-mediated guidance in neural crest cells in *X. laevis* via cellular repulsion (Smith et al., 1997). However, re-using developmental guidance cues in a mature adult environment following injury may not be effective since the location of the injury and the tissues needing replacement can vary unpredictably whereas developmental events are orderly and synchronized (Guedelhoefer IV. & Alvarado, 2012; Laird et al., 2008). If irrefutable evidence for migration of Vasa-positive coelomic cells is found in *C. teleta*, next it would be valuable to identify what guidance cues these cells use to navigate to the posterior wound.

### Activation of Wnt/β-catenin Signaling Does Not Lead to Regeneration Completion

In a previous study, we elicited partial posterior regeneration by increasing Wnt/β-catenin signaling in tail fragments cut between segment 10 and 11 (Kunselman & Seaver, 2025). We therefore wondered if the increased regenerative potential in 1-2 amputees relative to 10-11 amputees could lead to complete ectopic posterior regeneration upon activation of Wnt/β-catenin signaling. We find that 1-2 amputees treated with CHIR either from 0-3 dpa or continuously for 7 dpa do not progress to further steps of regeneration than do 10-11 amputees. Originally, we thought this could be because additional organizing signals downstream of canonical Wnt signaling are necessary for later stages of regeneration such as segment formation. However, we observed two 10-11 amputees that showed signs of complete regeneration after treatment with CHIR. These individuals prove that it is possible, although rare, to achieve complete posterior regeneration by activating Wnt/β-catenin signaling alone. A possible reason that 1-2 amputees do not complete posterior regeneration at a higher frequency than 10-11 amputees could be because tissue with anterior identity is refractory to transforming into tissue with posterior identity. This hypothesis could be tested by applying CHIR to tail fragments amputated several segments posterior to segment 10 and observing if bicaudal animals form at a higher frequency from the more posterior amputation site. Another potential reason 1-2 amputees fail to regenerate a complete tail after CHIR treatment could be because a signaling pathway is activated in the anterior-facing blastema that is antagonistic to posterior regeneration. In the future, comparative transcriptomics of the anterior- and posterior-facing blastemas of *C. teleta* can serve as an unbiased way to screen for important signaling pathways that may be influencing regeneration outcome and identity.

### Differences Between Anterior-facing and Posterior-facing Blastemas

As stated earlier, we currently cannot be certain of the identity of the anterior-facing blastema of 1-2 amputees. Since no differentiated structures specific to a head or tail appear in the regenerate, further characterization of anterior- or posterior-marker genes may be necessary to reveal the polarity of the blastema. There are few annelid studies that both identify genes uniquely expressed in the anterior-facing blastema and verify them *in situ* (Bely & Wray, 2001; Takeo et al., 2008). However, RNAseq studies comparing gene expression differences between the anterior and posterior blastema of annelids that regenerate in both directions could supply some candidate orthologs that may be exclusively expressed in the anterior-facing blastema of *C. teleta* (Bely & Wray, 2001; Ribeiro et al., 2019). Since anterior regeneration is believed to have been present in the last common ancestor of annelids, *C. teleta* may have retained some remnant of an ancestral anterior regeneration gene regulatory network (Zattara & Bely, 2016).

Notably, the anterior-facing blastema in *C. teleta* displays different characteristics than posterior-facing blastemas that regenerate a tail: 1) it does not co-express *Ct-piwi1* and *CapI-vasa* at 3 dpa, 2) its regeneration ability is not enhanced by or reliant on Vasa-positive coelomic cells, and 3) it has different domains of Pax expression, serotonin-positive neurites, and FMRFamide-positive neurites relative to regenerating posterior tissue (for posterior regeneration patterns, see Fig. S5). Further analyses could identify additional differences between the anterior- and posterior-facing blastemas of *C. teleta* and lead to better understanding of the distinct and shared mechanisms between anterior and posterior regeneration. Then, these data could help explain the evolutionary patterns of frequent loss of anterior regeneration ability despite mostly preserved posterior regeneration among the annelids (Zattara & Bely, 2016).

### Summary and Conclusion

We demonstrate that tail fragments cut between segments 1 and 2 have higher regeneration potential than tail fragments cut between segments 10 and 11. The increased regeneration potential of tail fragments cut between segments 1 and 2 is not due to the presence of the MPC cluster, a putative source of somatic stem cells in *C. teleta*. Activation of Wnt/β-catenin signaling at the more anterior cut site does not lead to complete posterior regeneration. Although anterior-facing and posterior-facing blastemas share some gene expression patterns and cellular responses following amputation, differences between them suggest that they form via distinct mechanisms.

Our study reveals that regeneration research in *C. teleta* can contribute to many major unsolved mysteries of regenerative biology. *C. teleta* forms an anterior-facing blastema seemingly via distinct mechanisms and cell sources from the posterior blastema, so within one organism, two mechanisms of blastema formation can be compared. The underlying causes for the variation of regenerative ability along the AP axis of *C. teleta* also remain to be determined, and by doing so *C. teleta* can become a valuable point of comparison for other animals with limited regeneration potential. Understanding the regulatory events driving tissue regeneration can provide the means to rescue latent regeneration abilities, and annelids have been understudied in this regard thus far. The diversity of regeneration mechanisms among annelids is exciting because comparative regeneration studies within this taxon can elucidate evolutionary patterns of regeneration mechanisms that are difficult to extrapolate from broad comparisons across multiple phyla (Bely, 2006). Comparisons of cell behavior during annelid regeneration can reveal the guiding principles of both stem-cell based and stem-cell independent regeneration, which may apply to other organisms. Annelid regeneration research is poised to make revolutionary contributions to the regeneration biology field.

## Acknowledgments

Research support came from The Carl and Marcella Matthaei Ecological Scholarship Fund to LFK and the National Science Foundation to ECS (IOS 2316882).

