## Supplemental Figures for "Regenerative potential varies along the anterior-posterior axis of the annelid *Capitella teleta*"

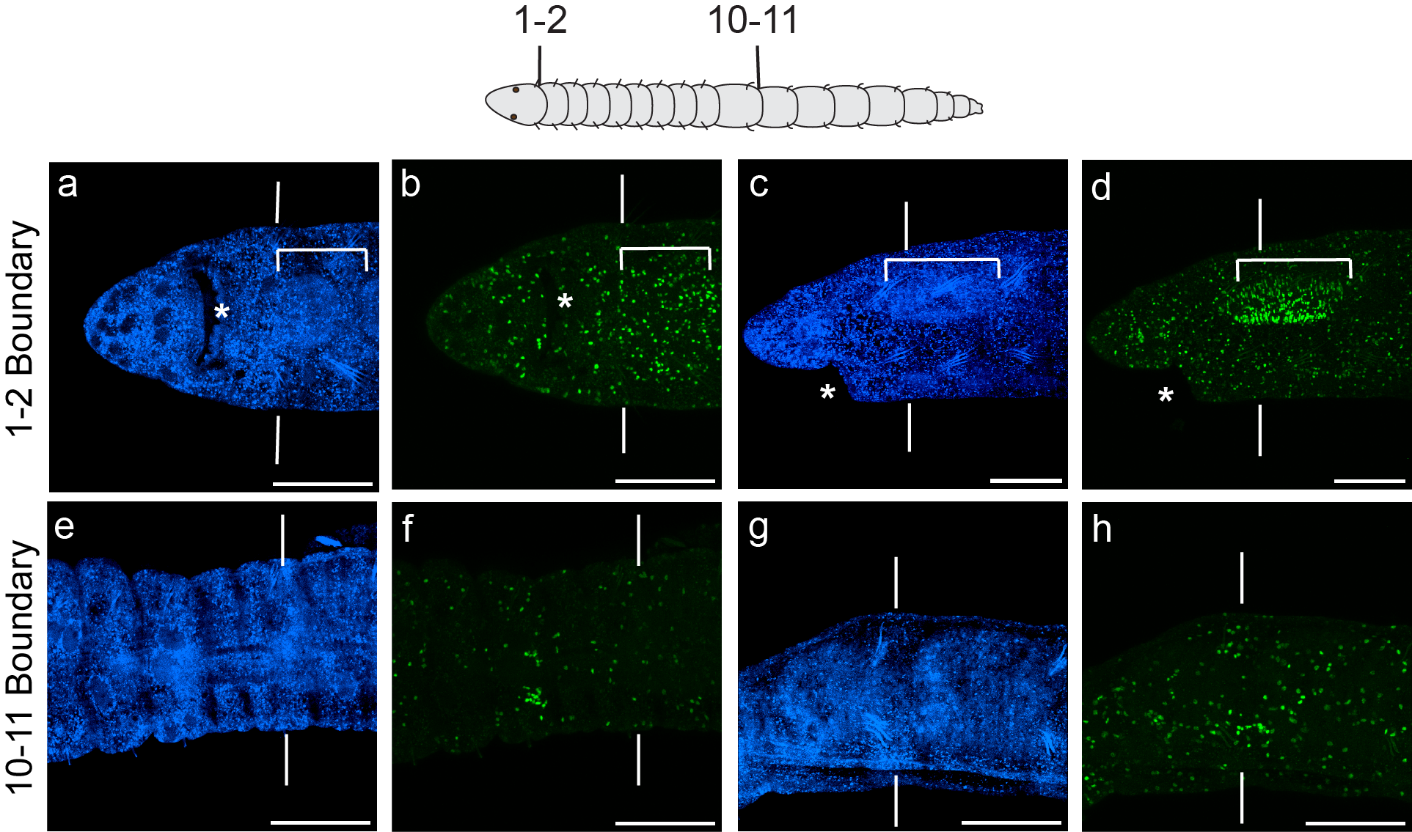

Figure S1. Intact animals have distinct patterns of EdU incorporation along the AP axis. a-d) The tissue surrounding the boundary between segments 1 and 2. e-h) The tissue surrounding the boundary between segments 10 and 11. Nuclei stained with Hoechst 33342 are blue and EdU+ nuclei are green. Asterisk indicates the mouth. Vertical lines indicate segment boundaries. Brackets encompass the pharynx. In all images, anterior is to the left. In leftmost two columns, view is ventral. In righthand two columns, view is lateral. Scale bars, 100 µm.

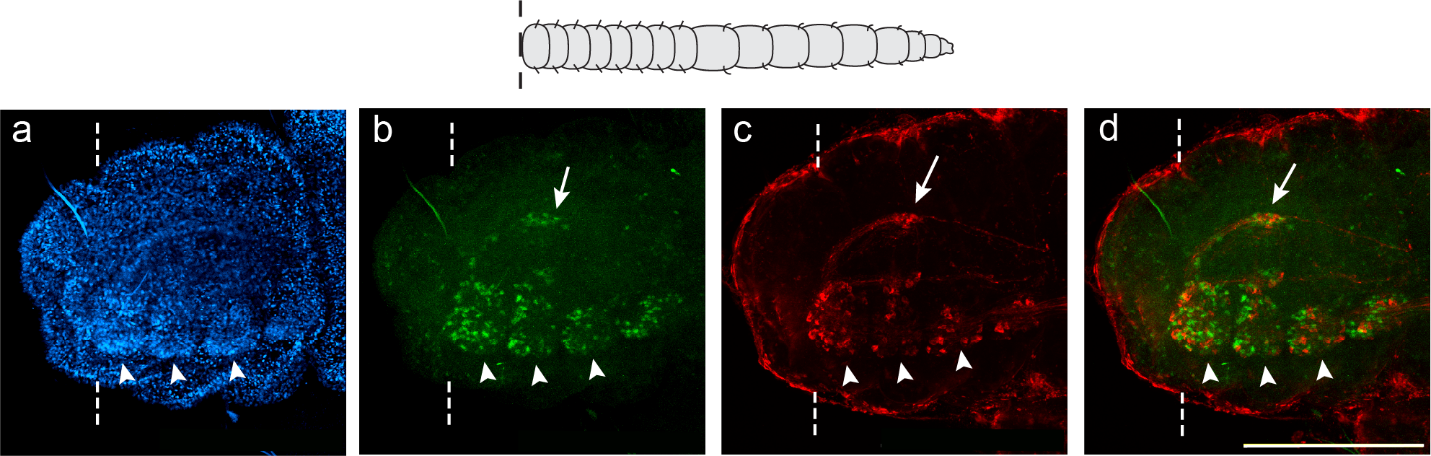

Figure S2. Dorsal domain of Pax expression colocalizes with FMRFamide nerve extensions. Panels depict a single tail fragment cut between segments 1 and 2 and allowed to regenerate for 7days. a) Nuclei in blue are stained with Hoechst 33342, b) Anti-Pax expression is in green, c) Anti-FMRFamide is in red. d) Merge of anti-Pax and anti-FMRFamide channels. Arrows point to dorsal Pax and FMRFamide expression. Arrowheads point to ganglia of the VNC. Dotted lines indicate amputation site. In all panels, anterior is to the left and ventral is down. Scale bar, 100 µm.

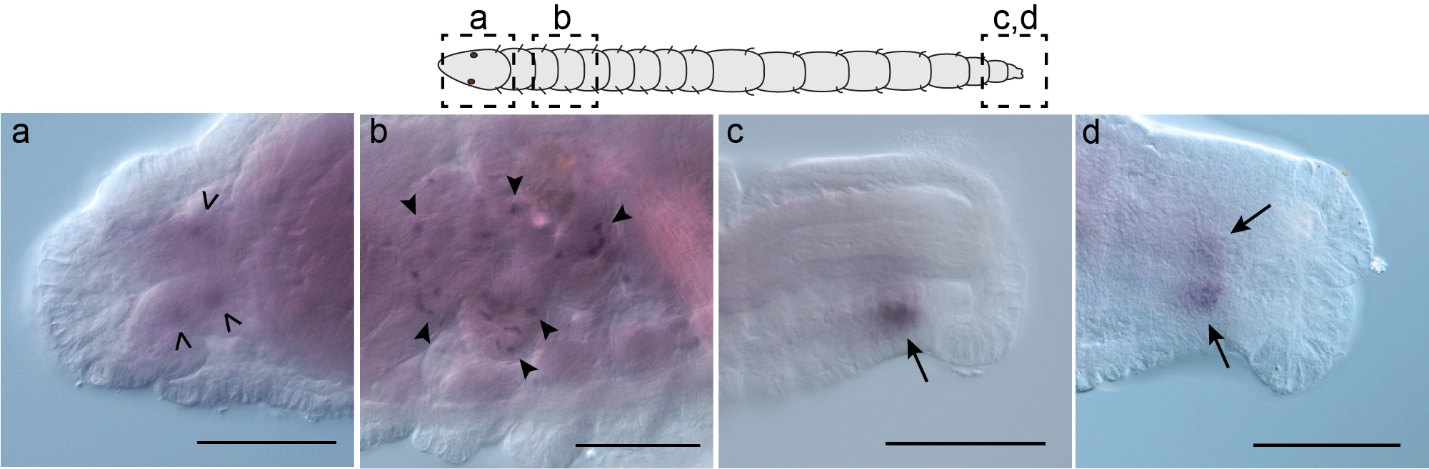

Figure S3. *Ct-neuroD* expression in intact *C. teleta* juveniles. a) Open arrowheads point to expression in the brain. b) Solid arrowheads point to expression in cells in the enteric nervous system posterior to the pharynx. c) Arrow points to expression in the pgz from a lateral view. d) Arrows point to bilateral expression in the pgz from a ventral view. In all images, anterior is to the left. Scale bar 100 µm.

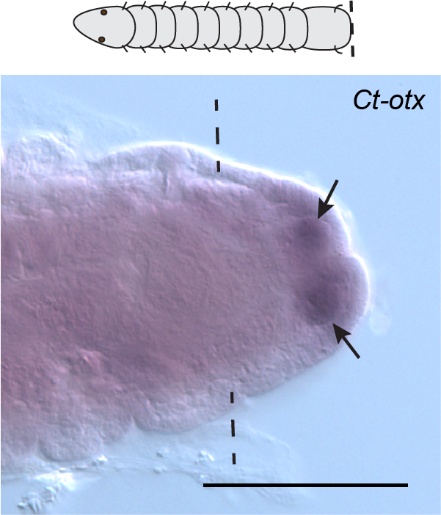

Figure S4. *Ct-otx* is expressed in the mesoderm of the blastema of regenerating head fragments 3 dpa. Arrows point to bilateral expression of *Ct-otx* in the mesoderm of the blastema. Amputation site is posterior to segment 10. Dotted lines indicate amputation site. View is ventral. Anterior is to the left. Scale bar 100 µm.

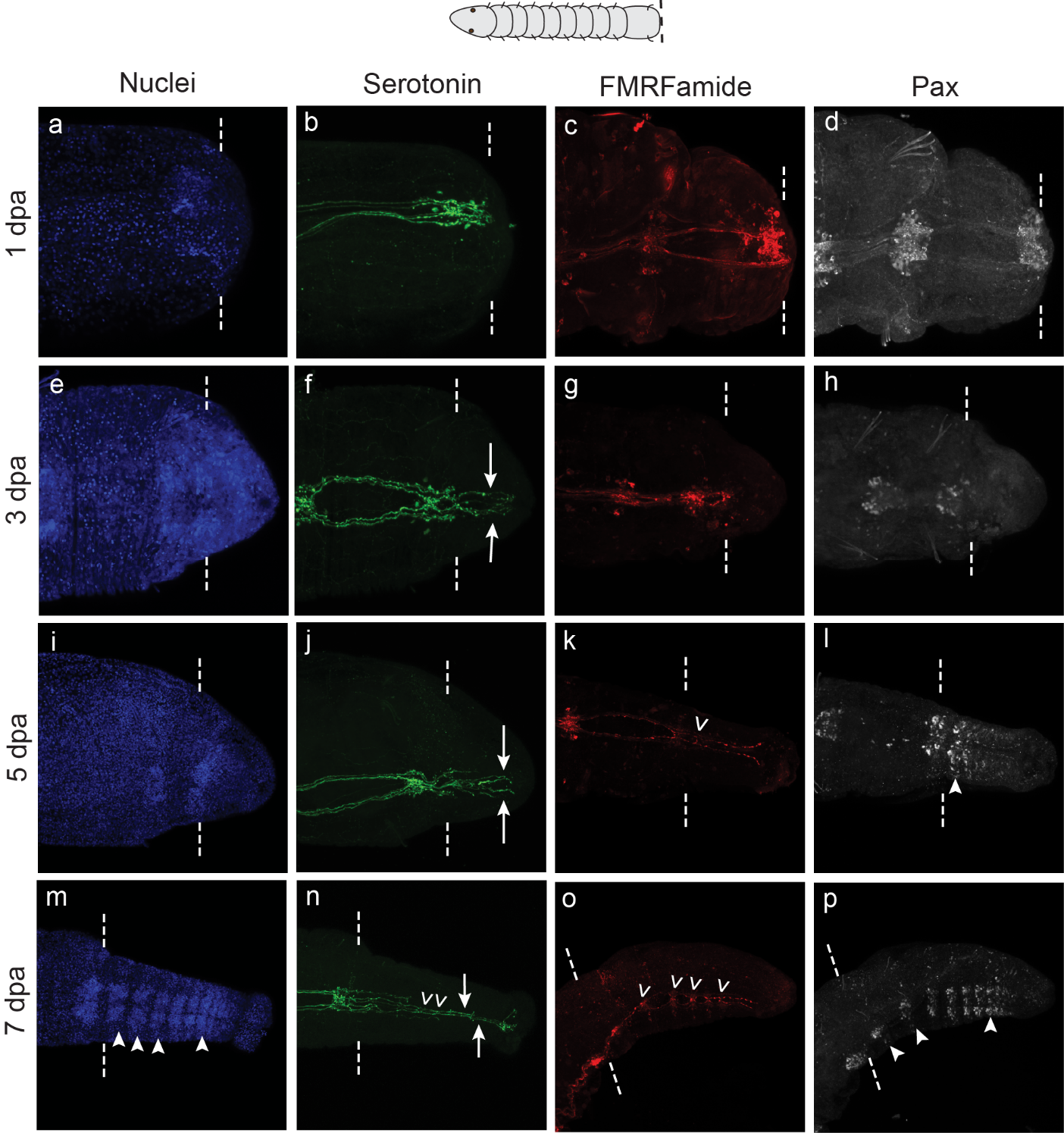

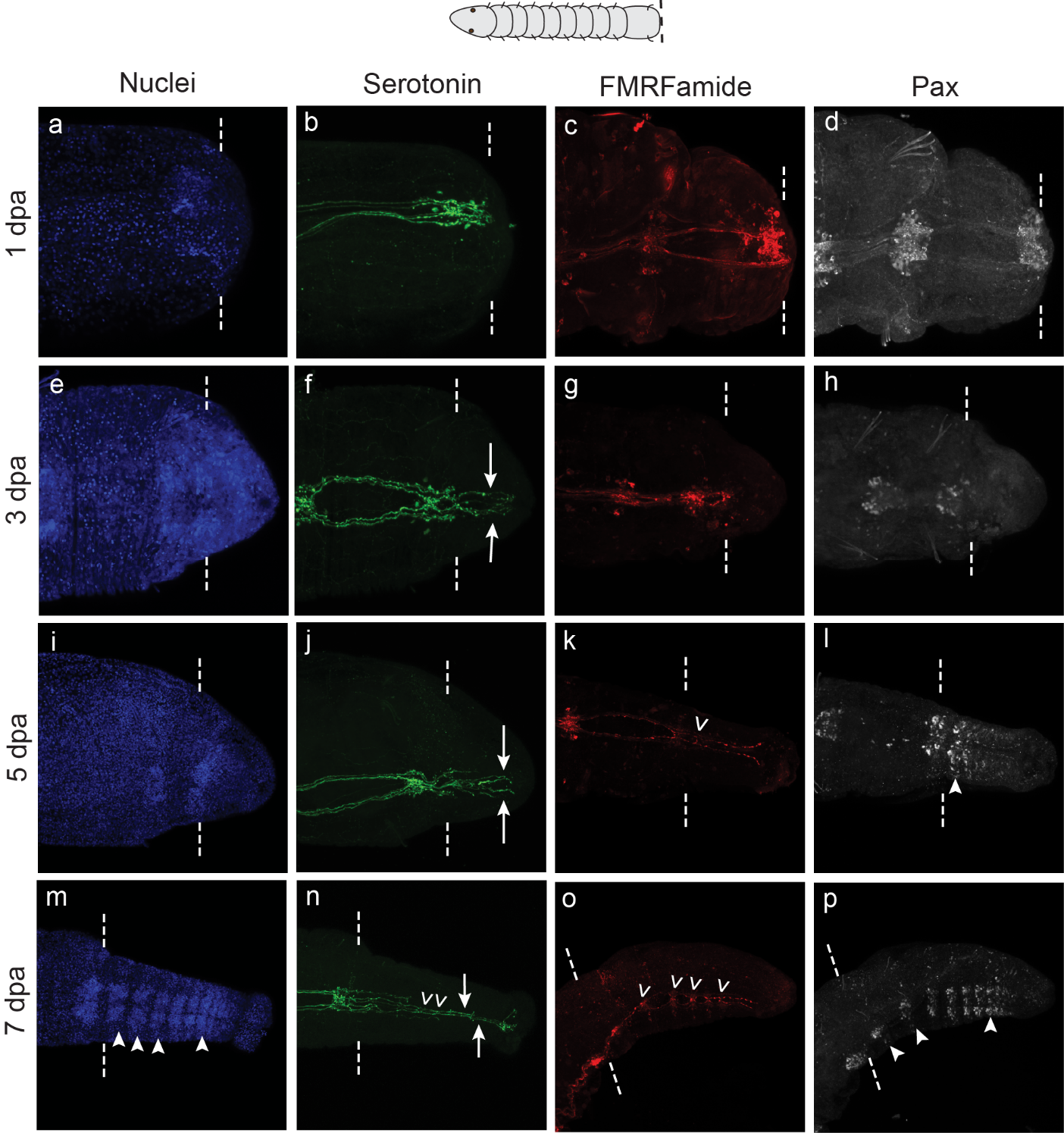

Figure S5. Patterns of neural subtypes in posterior regeneration. Head fragments were cut posterior to segment 10 and fixed 1 dpa (a-d), 3 dpa (e-h), 5 dpa (i-l), or 7 dpa (m-p). Nuclei are stained blue with Hoechst 33342, anti-serotonin is green, anti-FMRFamide is red and anti-Pax is white. Arrows point to extensions of ventromedian serotonin+ connectives. Open arrowheads point to commissures between longitudinal neurites. Solid arrowheads point to new ganglia. Dotted lines indicate amputation site. In the left-hand two columns, panels for each time point are from the same animal. In the right-hand two columns, panels for each time point are from the same animal. All images are in ventral view and anterior is to the left.
